# Phosphoproteomics reveals that the hVPS34 regulated SGK3 kinase specifically phosphorylates endosomal proteins including Syntaxin-7, Syntaxin-12, RFIP4 and WDR44

**DOI:** 10.1101/741652

**Authors:** Nazma Malik, Raja S Nirujogi, Julien Peltier, Thomas Macartney, Melanie Wightman, Alan R Prescott, Robert Gourlay, Matthias Trost, Dario R. Alessi, Athanasios Karapetsas

## Abstract

The serum- and glucocorticoid-regulated kinase (SGK) isoforms contribute resistance to cancer therapies targeting the PI3K pathway. SGKs are homologous to Akt and these kinases display overlapping specificity and phosphorylate several substrates at the same residues, such as TSC2 to promote tumor growth by switching on the mTORC1 pathway. The SGK3 isoform is upregulated in breast cancer cells treated with PI3K or Akt inhibitors and recruited and activated at endosomes, through its phox homology domain binding to PtdIns(3)P. We undertook genetic and pharmacological phosphoproteomic screens to uncover novel SGK3 substrates. We identified 40 potential novel SGK3 substrates, including 4 endosomal proteins STX7 (Ser126) and STX12 (Ser139), RFIP4 (Ser527) and WDR44 (Ser346) that were efficiently phosphorylated *in vitro* by SGK3 at the sites identified *in vivo,* but poorly by Akt. We demonstrate that these substrates are poorly phosphorylated by Akt as they possess an n+1 residue from the phosphorylation site that is unfavorable for Akt phosphorylation. Phos-tag analysis revealed that stimulation of HEK293 cells with IGF1 to activate SGK3, promoted phosphorylation of a significant fraction of endogenous STX7 and STX12, in a manner that was blocked by knock-out of SGK3 or treatment with 14H inhibitor. SGK3 phosphorylation of STX12 enhanced interaction with the VAMP4/VTI1A/STX6 containing SNARE complex and promoted plasma membrane localization. Our data reveal novel substrates for SGK3 and suggest a mechanism by which STX7 and STX12 SNARE complexes are regulated by SGK3. They reveal new biomarkers for monitoring SGK3 pathway activity.

## Introduction

The phosphoinositide 3-kinase (PI3K) pathway plays crucial role in various biological processes, including cell growth, proliferation and metabolism [1]. Most types of tumors harbor mutations that hyperactivate the PI3K pathway and a plethora of inhibitors are being evaluated as anticancer agents [2]. Recently an orally available PI3K alpha specific inhibitor termed Alpelisib has been approved for treatment of PIK3CA-mutated, hormone receptor-positive advanced breast cancer [3]. Akt inhibitors Capivasertib [4] and Ipatasertib [5] are in late stage clinical trials for breast cancers bearing PI3K or Akt mutations. However, studies to date indicate that despite encouraging initial responses, the majority of tumors display inherent resistance or develop acquired resistance to PI3K/Akt pathway inhibitors through diverse mechanisms [6, 7]. One of these resistance pathways involves upregulation of the family of serum- and glucocorticoid-regulated kinases, (SGKs), which belong to the AGC family of kinases and bear high similarity to Akt, especially in the catalytic domain. There are 3 isoforms of SGK termed SGK1, SGK2 and SGK3 [8, 9]. Breast cancer cell lines carrying mutations in PTEN or PI3K that also express high levels of SGK1, display inherent resistance to PI3K or Akt inhibitors [10]. Moreover, clinical trials with PI3Kα inhibitors reveal that elevation of SGK1 activity contributes to therapy resistance, at least in part by phosphorylating TSC2 and activating the mTORC1 pathway [11]. In breast cancer cell lines that are initially sensitive to PI3K or Akt inhibitors and that express low levels of SGK1, prolonged treatment with PI3K or Akt inhibitors resulted in upregulation and activation of SGK3 that also substituted for Akt by phosphorylating TSC2 and ultimately activating mTORC1 [12].

Similar to Akt isoforms, the 3 isoforms of SGK are activated by phosphorylation of their kinase domain T-loop (Thr320 residue in SGK3) by PDK1 (3-phosphoinositide dependent kinase-1) and at a C-terminal hydrophobic motif (Ser486 in SGK3) by mTORC2 (mTOR complex 2) [13-17]. Phosphorylation of the hydrophobic motif by mTORC2 creates a docking site for PDK1 to phosphorylate the T-loop residue resulting in the activation of SGK isoforms [15-18]. SGK1 and SGK2 unlike Akt isoforms do not possess a regulatory phosphoinositide (PtdIns) binding domain and their activity is therefore tightly controlled by the activity of mTORC2 [9]. SGK3 possesses an N-terminal PtdIns(3)P-binding PX (Phox homology) domain [19]. This targets SGK3 to endosomal membranes, where a large pool of PtdIns(3)P is generated by the Class III PI3K family member, known as hVps34 [17, 19, 20]. Activation of Class I PI3K can also contribute to the pool of PtdIns(3)P at the endosomal membrane through the sequential degradation of PtdIns(3,4,5)P3 via SHIP2 and INPP4A/4B inositol phosphatases [21, 22]. Studies undertaken with selective class 1 and class 3 PI3K inhibitors indicate that both PI3K family members contribute to the activation of SGK3 by growth factors [22].

SGK isoforms phosphorylate substrates in a RXRXXS/T motif similar to Akt [13, 23, 24]. Indeed, Akt and SGK isoforms phosphorylate many substrates, including TSC2 [11, 12], Bad [25], NEDD4L [26-28] and FKHRL-1 [29] at the same residues. The most commonly studied *in vivo* readout of SGK isoform activity is NDRG1 (N-Myc downstream-regulated gene 1), which is efficiently phosphorylated at Thr346 by Akt [30], SGK1 [10] as well as SGK3 [12, 22]. Only two SGK3 substrates have been reported namely AIP4 [31] and FLI-1 [32] that were apparently not phosphorylated by SGK1 and SGK2. To our knowledge these substrates have not been independently confirmed by others and it is not known whether these proteins are phosphorylated by Akt.

Akt has a strong preference for a large hydrophobic residue such as Phe at the n +1 residue [23]. A substrate peptide for SGK1 termed MURRAYtide derived from NDRG1 was elaborated in previous work that was poorly phosphorylated by Akt [33]. The MURRAYtide peptide possesses a serine residue at the n+1 position, and changing this to a Phe residue markedly improved phosphorylation by Akt [33]. This work suggested that the n+1 position might represent an important determinant of whether a substrate is phosphorylated by Akt. A prediction from this work is that SGK selective substrates not phosphorylated by Akt, possess non-large hydrophobic residues at the n+1 position. However, the Thr346 residue of NDRG1 that is phosphorylated by Akt and SGK isoforms *in vivo* [10], lies within the RSRSHpTS sequence motif and therefore has a Ser residue as the n+1 residue. This demonstrates that at least in the context of a protein, a substrate bearing a non-optimal n+1 residues can be phosphorylated by Akt *in vivo*. In this study, we set out to identify whether there are any SGK3 selective substrates that were not phosphorylated by Akt and study the roles that these might play.

## Results

### Phosphoproteomic Screens to identify novel SGK3 substrates

To identify physiological substrates of SGK3, we compared the phosphoproteomes of wild type (n=4) and SGK3 knockout (n=4) serum starved HEK293 cells treated with an Akt inhibitor (1 µM MK2206) prior to stimulation with IGF1 (50 ng/ml, 15 min). This phosphoproteome screen was termed PS1 (Fig 1A left panel). Control immunoblotting analysis confirmed that the dual Akt/SGK NDRG1 substrate, was phosphorylated to a much greater extent in the wild type compared to SGK3 knock-out cells (Fig 1A left panel). The residual phosphorylation of NDRG1 in SGK3 knock-out cells is likely due to low levels of SGK1 activity [22]. We also performed a second screen termed PS2, in which we compared the phosphoproteomes of IGF1-stimulated (50 ng/ml, 15 min), serum-starved wild type HEK293 cells treated with either no inhibitor (DMSO control n=3), with a pan SGK isoform inhibitor (3 µM 14H, 1 h, n=3) or with an Akt inhibitor (1 µM MK2206, 1 h, n=3) (Fig 1A right panel). Control immunoblotting revealed that phosphorylation of NDRG1 was partially reduced in SGK inhibitor (14H) or Akt inhibitor (MKK2206) treated samples (Fig 1A right panel), similar to what has been observed in a previous study [22]. Phosphorylation of the Akt-selective substrate PRAS40 that is not phosphorylated by SGK isoforms, was ablated by the Akt inhibitor (MK2206) but not the SGK inhibitor (14H) treatment (Fig 1A).

**Figure 1:**
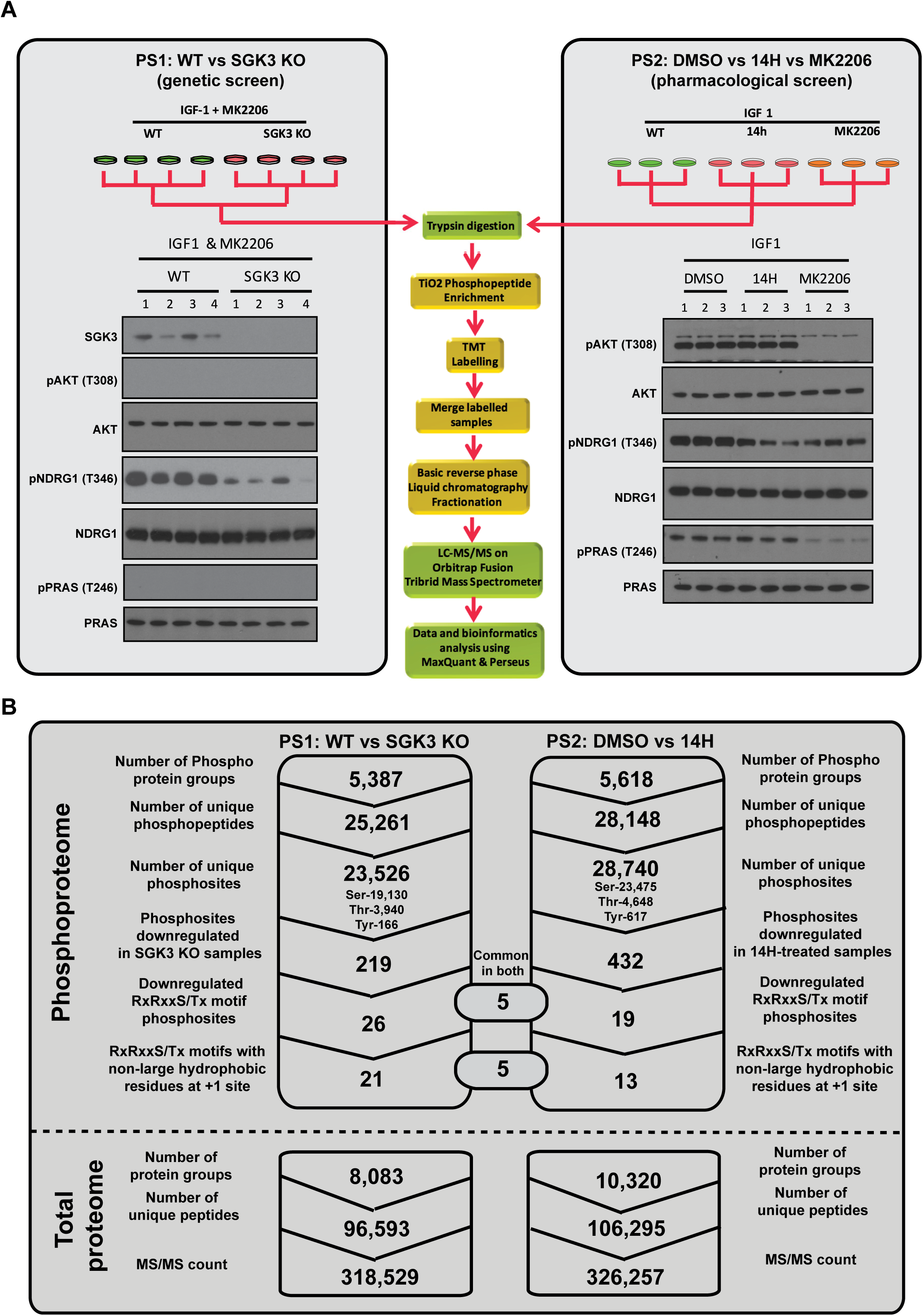
Quantitative phosphoproteomics using TMT (Tandem mass tag) labelling to identify selective SGK3 substrates. A) TMT Phosphoproteomics workflow. Wild type (WT) and SGK3 knock-out (KO) HEK293 cells were serum starved overnight, treated then with 1µM MK2206 for 3 h and then stimulated with 50 ng/ml IGF1 for 15 min (PS1, left panel) or WT HEK293 cells were serum starved overnight, treated for 1 h with 1µM MK2206, 3µM 14H or left untreated (DMSO) and then stimulated with 50 ng/ml IGF1 (PS2, right panel). Cells were lysed in 2% by mass SDS lysis buffer and control immunoblotting was performed with the indicated antibodies. For the TMT labelling, after lysis samples were trypsinized and then phosphopeptide enrichment was performed using TiO_2_ beads after which the peptides were labelled with TMT and mixed (for PS1 127N, 127C, 128N and 128C were labelled to WT-HEK293 samples and 129N, 129C, 130N and 130C were labelled to SGK3-KO samples; for PS2 the TMT reporter tags, 126, 127N and 127C labelled the DMSO treated samples 128N, 128C and 129N labelled the MK2206 treated samples, 129C, 130N and 130C labelled the 14H treated samples). The TMT labelled samples were subjected to basic pH reverse liquid chromatography for fractionation and analyzed on Orbitrap Fusion Tribrid Mass spectrometer. B) Summary of the results depicting the identification of unique number of phosphosites and the total number of protein groups from phosphoproteome and total proteome analysis of PS1 and PS2 that were processed through MaxQuant pipeline.

Phosphoproteomic analysis was undertaken using a multiplexed quantitative phosphopeptide workflow outlined in Fig 1A, in which tryptic peptides from each sample were labelled with different isotopically labelled isobaric Tandem Mass Tags (TMT) and then mixed together prior to mass spectrometry analysis [34]. Mass spectrometry data was analyzed using the MaxQuant environment [35, 36]. All the mass spectrometry data for these screens has been deposited on the PRIDE database [37] with the accession number PXD014561. For PS1 we quantified >25,000 phosphopeptides from 5,387 protein groups and >28,000 phosphopetides from 5,618 protein groups in PS2 (Fig 1B). For PS1, 219 phosphorylation sites were significantly reduced in all 4 replicates of the SGK3 knock-out samples. For PS2, 432 phosphorylation sites were significantly reduced in all 3 replicates of SGK inhibitor (14H) treated samples (Fig 1B). The independently acquired total proteome measurements confirmed that the detected phosphorylation changes in PS1 and PS2 were not due to altered protein abundances (changes as determined by MaxQuant were less than twofold [Fig S1]).

The high confidence phosphosites impacted by SGK3 knock-out in PS1 (Fig 2A) or inhibition of SGK3 in PS2 (Fig 2B) are illustrated by volcano plots. We focused our subsequent analysis on impacted phosphorylation sites lying within the well characterized Akt and SGK phosphorylation site consensus RXRXXS/T motif [13, 23], as these represent good candidates for direct SGK3 substrates. We identified 26 robust phosphorylation sites in PS1 reduced by SGK3 knock-out and 19 robust phosphorylation sites in PS2 inhibited with 14H, within RXRXXS/T motif (Fig 1B, Fig 2C). Five of these hits were independently identified in both PS1 and PS2. These included Neural Developmentally Down-Regulated 4-Like E3 ligase (NEDD4L, Ser385) that as discussed in the Introduction is a validated Akt, SGK1 and SGK3 substrate [27, 28], and four proteins that to our knowledge not previously described as SGK or Akt substrates, namely Syntaxin-7 (STX7, Ser126), Syntaxin-12 (STX12, Ser139), Myc-Binding Protein-2 E3 ligase (MYCBP2, Ser2833) and Intersectin-1 (ITSN1, Thr278). The proteomic screens also identified the previously characterized Akt and SGK NDRG1 substrate (Ser366, PS1) and (Ser326, PS2) as well as the related NDRG3 protein (Ser331, PS1). Three other hits corresponded to previously reported Akt substrates namely BRISC And BRCA1 A Complex Member-1 (BABAM1/ MERIT40) (Ser29, PS1) [38], Ataxin-1 (Ser775, PS1) [39] and WDR44 (Ser344, PS1) [40]. The evidence that WDR44 is phosphorylated by Akt is based on biochemical analysis showing that immunoprecipitated Akt phosphorylated WDR44 at Ser342 and Ser344 [40]. However, mass spectrometry analysis we have undertaken suggest Ser344 which lies in an RXRXXS/T motif is the likely site of phosphorylation (>95% confidence) (Fig 2C, Supplementary Excel file). The Ser342 residue does not lie within an Akt/SGK phosphorylation motif.

**Figure 2:**
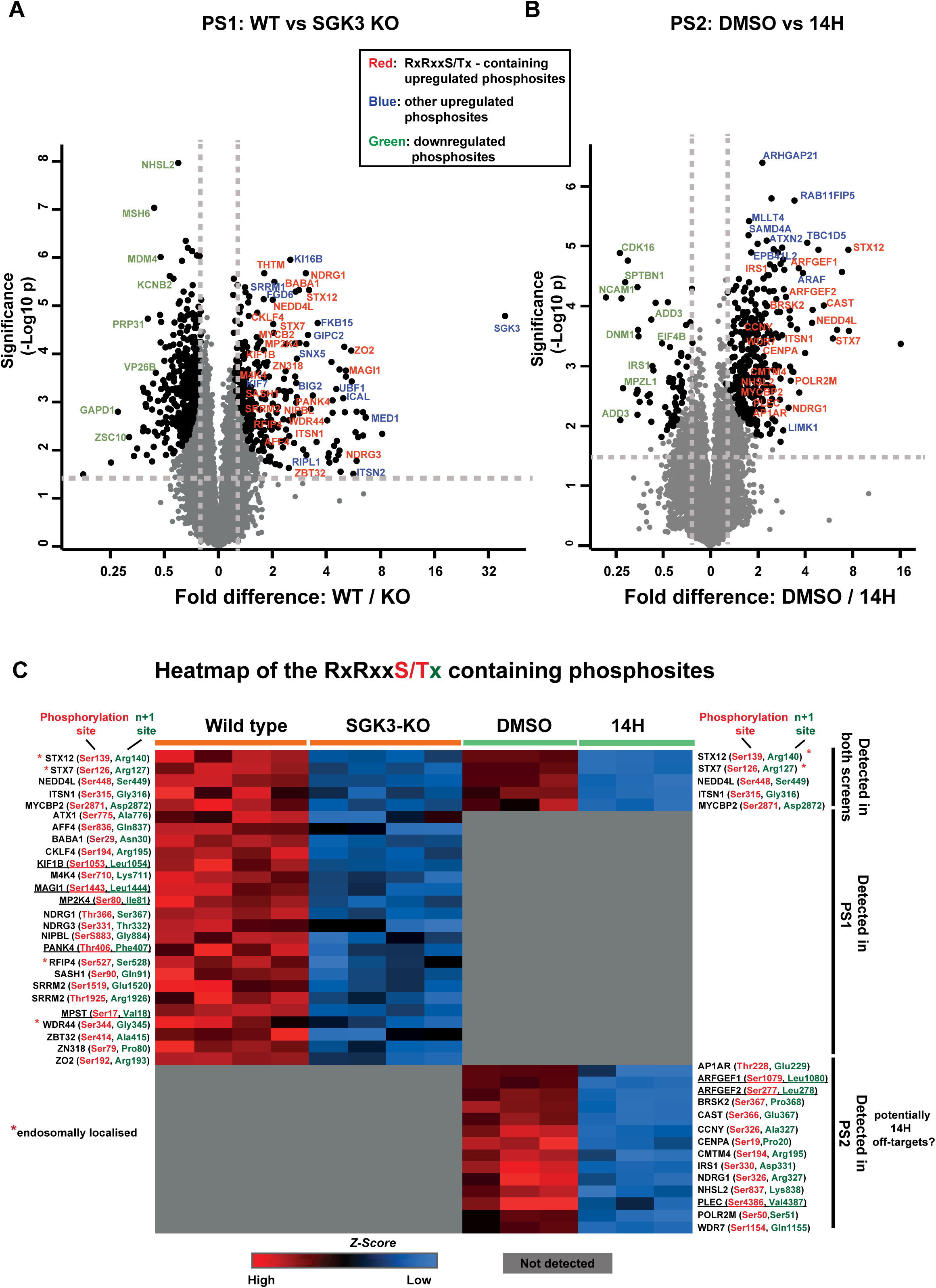
Identification of novel SGK3 selective substrates. Volcano plots depicting the phosphoproteome statistical significance between the HEK293 Wild type (WT) and SGK3 knock-out (KO) cells (N=4) from PS1 on the left panel (A) and for the HEK293 cells treated with DMSO and 14H (N=3) from PS2 on the right panel (B). A) For PS1, the significantly enriched phosphosites in WT and KO conditions are highlighted in enlarged black circles along with the color coded text; In red fonts phosphosites that are specific to RXRXXS/T motif and enriched in the wild type samples, in blue other phosphosites enriched in the wild type samples and in green phosphosites enriched in the SGK3 KO samples. B) For PS2, the significantly enriched phosphosites in DMSO and 14H conditions are highlighted in enlarged black circles along with the color coded text; In red fonts phosphosites that are specific to RXRXXS/T motif and enriched in the DMSO treated samples, in blue other phosphosites enriched in DMSO treated samples and in green phosphosites enriched in 14H samples. For both screens the statistical significance was defined by the independent two-tailed t-test, which is corrected by permutation-based FDR of 5% using the Perseus software. C) Heat map depicting the summary of statistically significant RXRXXS/T motif containing phosphosites identified in the two screens. The log2 values of each of the RXRXXS/T phosphosite is Z-score normalized and generated the heat map using Perseus software. The phosphorylation site and the +1 site of each motif are highlighted in red and green respectively. Phosphosites containing a large hydrophobic residue at the +1 site are underlined.

As SGK3 is located on the endosome, we perused the list of potential substrates reported to reside on endosomes. This analysis revealed five proteins, namely Intersectin-1 (ITSN1, Ser315, PS1 + PS2, n+1 = Gly) [41], Syntaxin-7 (STX7, Ser126, PS1 + PS2, n+1 = Arg) [42], Syntaxin-12 (STX12, Ser 139, PS1 +PS2, n+1 = Arg) [43], Rab11 family binding protein 4 (RFIP4, Ser527, PS1, n+1 = Ser) [44], and WD Repeat Domain 44 (WDR44, Ser344, PS1, n+1 = Gly) [45].

### STX7, STX12, RFIP4 and WDR44 are phosphorylated efficiently by SGK3 and poorly by Akt *in vitro*

We next tested whether the potential substrates identified in PS1 and PS2 were phosphorylated by SGK3 and Akt *in vitro* using an *in vitro* ^32^P-γ-MgATP phosphorylation assay to monitor substrate phosphorylation (Fig 3 and Fig S2). We were able to express 10 of the hits obtained from PS1 and PS2 namely MPST, PANK4, ITSN1, ATXN1, AFF4, BABA1 STX7, STX12, WDR44 and RFIP4 as full-length proteins in *E. coli*, that included the five endosomal proteins. We employed a concentration of recombinant active SGK3 and Akt1 that phosphorylated NDRG1 to a similar level in a parallel reaction (Fig S2A). We observed that five of the substrates tested, namely MPST (n+1 Val Fig S2B), ATXN1 (n+1 Ala Fig S2D), PANK4 (n+1 Phe Fig S2E) and AFF4 (n + 1 Gln Fig S2F) and one of the endosomal substrates namely ITSN1 (n+1 Gly Fig S2C), were phosphorylated similarly by Akt and SGK3 *in vitro*, in a manner that was blocked by addition of an Akt (MK2206) or SGK (14H) inhibitor.

**Figure 3:**
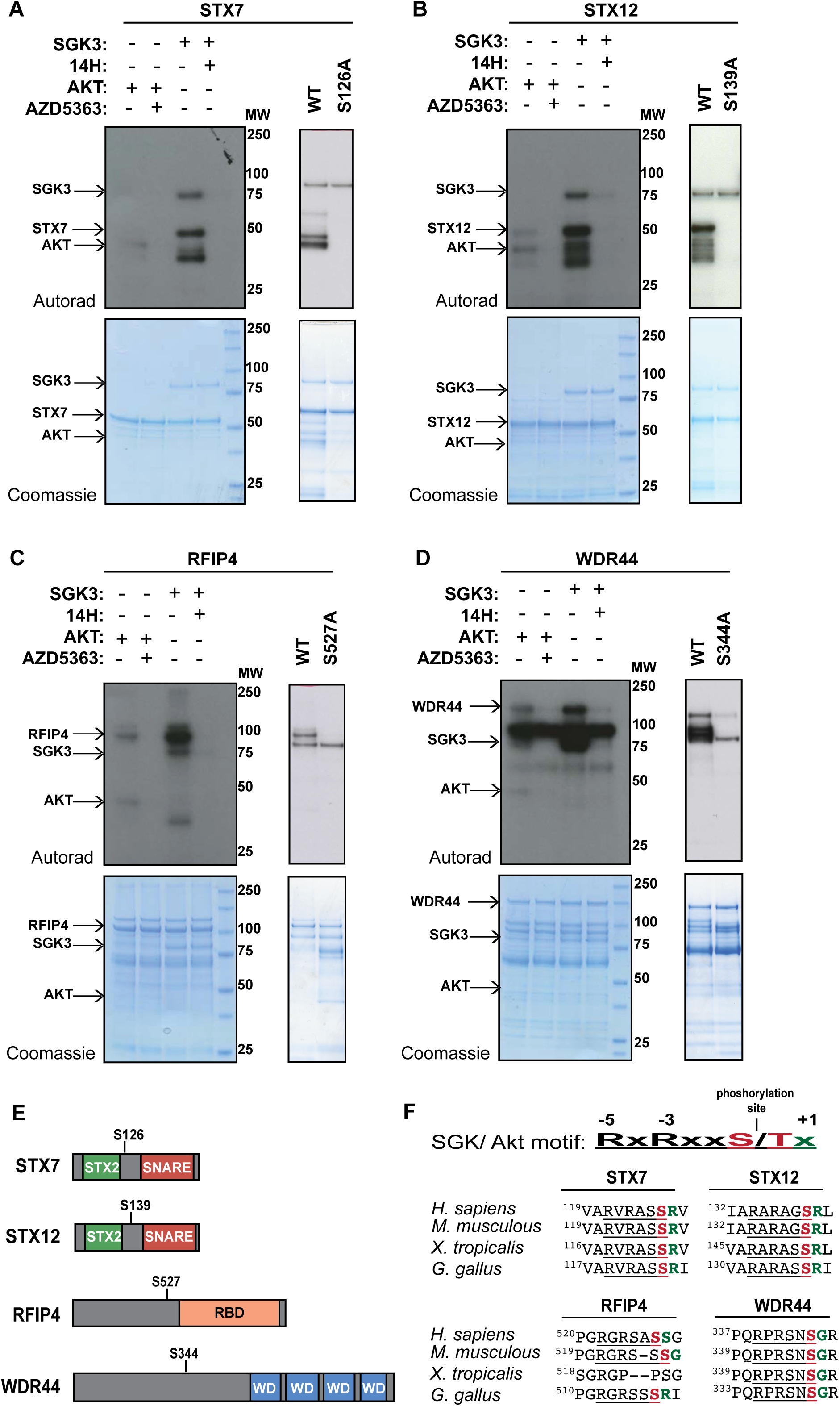
SGK3 but not Akt phosphorylates STX7, STX12, RFIP4 and WDR44 *in vitro*. 1-2 µg of GST-STX7 (A), GST-STX12 (B), GST-RFIP4 (C) and GST-WDR44 (D) were incubated with 0.5 µg active SGK3 or 0.125 µg active Akt and [γ^32^P]-ATP, in the presence or absence of 14H or AZD5363 (1 µM) for 30 min. The reactions were terminated by the addition of SDS loading buffer and separated by SDS/PAGE. Incorporation of [γ^32^P]-ATP was detected by autoradiography (top panels) and proteins were detected by Coomassie staining (bottom panels). Control kinase assays were also performed with the alanine mutants GST-STX7 S126A, GST-STX12 S139A, GST-RFIP4 S527A and GST-WDR44 S344A. E) Schematic representation of STX7, STX12, RFIP4 AND WDR44 domain organization. The location of the sites phosphorylated by SGK3 is also indicated. F) Sequence alignment of the motifs phosphorylated by SGK3 in STX7, STX12, RFIP4 and WDR44 in the indicated species. The alignment was performed using the Clustal omega tool (https://www.ebi.ac.uk/Tools/msa/clustalo/).

In contrast, the other four endosomal localized hits STX7 (n+1 Arg Fig 3A), STX12 (n+1 Arg Fig 3B), RFIP4 (n+1 Gly Fig 3C) and WDR44 (n+1 Ser Fig 3D) were phosphorylated by SGK3 to a significantly greater extent than Akt ranging from 3 to 10-fold higher levels of phosphorylation. Phosphorylation of STX7, STX12, RFIP4 and WDR44 by SGK3 was inhibited by an SGK inhibitor (14H) (Fig 3). Mutation of the site identified in the phosphoproteomic screens (STX7-Ser126, STX12-Ser139, RFIP4-Ser527 and WDR44-Ser344) to Ala, markedly inhibited the phosphorylation by SGK3 of all four substrates (Fig 3). We also performed *in vitro* ^32^P-label phospho-mapping analysis that confirmed that SGK3 specifically phosphorylates STX7 on Ser126 and RFIP4 on Ser527 (Fig S3). The domain structures of Syntaxin-7, Syntaxin-12, Rab11FIP4 and WDR44 and location of the SGK3 phosphorylation sites are summarized in Fig 3E.

### Mutation of n+1 residue to Phe enhances phosphorylation of STX7, STX12 and RFIP4 by Akt without impacting phosphorylation by SGK3

As mentioned above STX7, STX12, RFIP4 and WDR44 all possess an unfavorable n+1 residue that might account for why these proteins are poorly phosphorylated by Akt (Fig 3F). To address this question, we mutated the n+1 residue of STX7 (Fig 4A), STX12 (Fig 4B) and RFIP4 (Fig 4C) to a Phe residue that is favorable for phosphorylation by Akt. The recombinant proteins with a n+1 Phe were phosphorylated to a much greater extent than the wild type proteins by Akt (Fig 4). In contrast, the n+1 Phe mutation did not impact on phosphorylation of STX7, STX12 and RFIP4 by SGK3 (Fig 4). Using a concentration of Akt and SGK3 that phosphorylated NDRG1 similarly (Fig 4D), the STX7, STX12 and RFIP4 mutants possessing an n+1 Phe residue were similarly phosphorylated by both Akt and SGK3.

**Figure 4:**
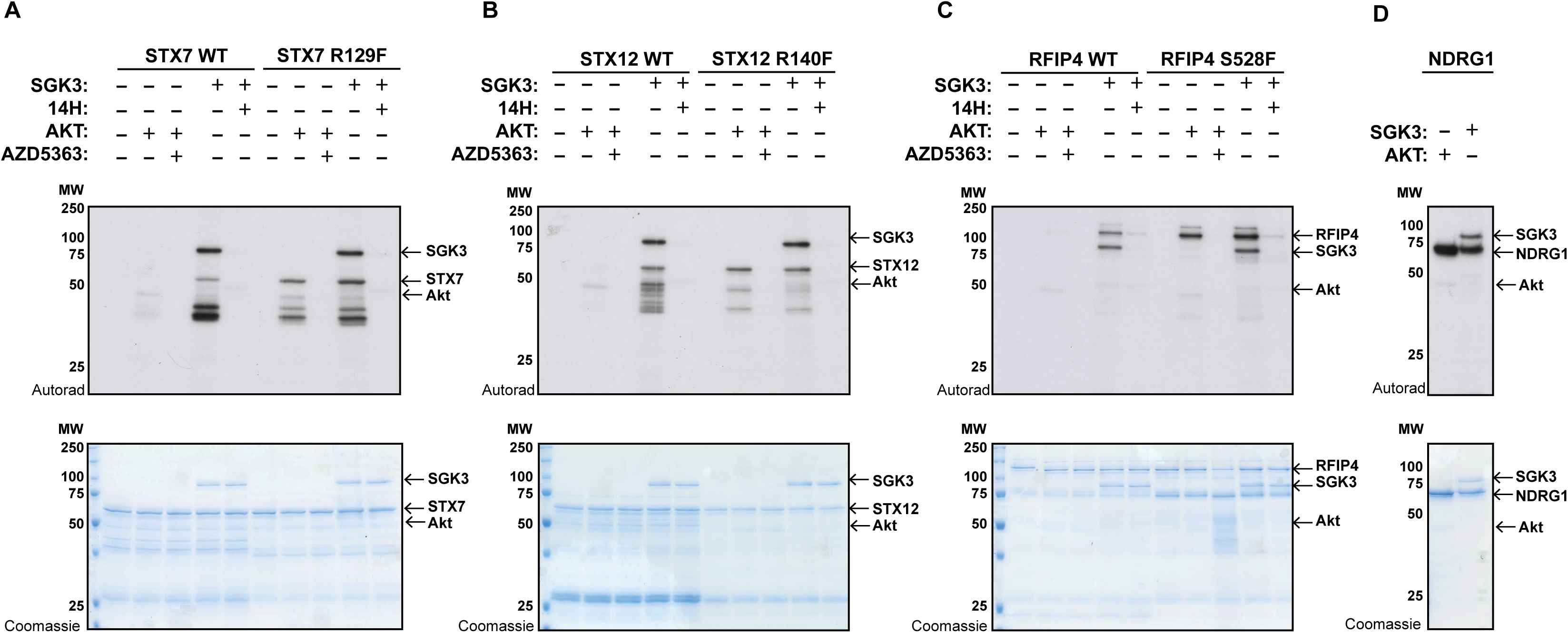
Mutation of n+1 residue to Phe enhances phosphorylation of STX7, STX12 and RFIP4 by Akt. 1-2 µg of GST-STX7 WT,GST-STX7 R127F (A), GST-STX12 WT, GST-STX12 R139F (B), GST-RFIP4 WT, GST-RFIP4 S528F (C) and GST-NDRG1 (D) were incubated with 0.5 µg active SGK3 or 0.1 µg active Akt and [γ^32^P]-ATP, in the presence or absence of 14H or AZD5363 (1 µM) for 30 min. The reactions were terminated by the addition of SDS loading buffer and separated by SDS/PAGE. Incorporation of [γ^32^P]-ATP was detected by autoradiography (top panels) and proteins were detected by Coomassie staining (bottom panels).

### STX7, and STX12 are efficiently phosphorylated by SGK1 *in vitro*

We next investigated whether STX7, STX12, RFIP4 and WDR44 could be phosphorylated by SGK1 similarly to SGK3 using *in vitro* ^32^P-γ-MgATP phosphorylation assay (Fig S4). We utilized a concentration of recombinant active SGK1, SGK3 and Akt1 that phosphorylated NDRG1 to similar levels in parallel reactions (Fig S4E). STX7 and STX12 were phosphorylated similarly by SGK1 and SGK3 (Fig S4A and S4B). WDR44 (Fig S4C) and RFIP4 (Fig S4D) were phosphorylated by SGK1 to a lesser extent compared to SGK3).

### Endogenous STX7 and STX12 are phosphorylated by SGK3 in HEK293 cells

We next explored whether the method known as “phos-tag gel electrophoresis” that retards the electrophoretic mobility of phosphorylated proteins [46, 47], was suitable for analyzing SGK3 mediated phosphorylation of endogenous STX7 and STX12 *in vivo*. We stimulated serum starved HEK293 cells ± IGF1 (15 min, 50 ng/ml) to activate SGK3 and immunoblotted for total STX7 and STX12 proteins after phos-tag gel electrophoresis. In serum-starved cells STX7 migrated as a single species, but following IGF1 stimulation a striking band-shift was observed indicating that a substantial fraction corresponding to ∼50% of the protein was phosphorylated (Fig 5A). The band-shifted species of STX7 migrated as a major and minor band suggesting that there is phosphorylation at a second minor site. Consistent with phosphorylation being mediated by SGK3, the appearance of both upper bands was blocked with the SGK inhibitor (14H 1 µM), but phosphorylation was not impacted by Akt inhibitor treatment (MK2206, 1 µM) (Fig 5A). Additionally, the IGF1-induced band-shift was blocked in SGK3 knock-out cells (Fig 5B). Control immunoblotting confirmed that IGF1 activated Akt and SGK3 and that the inhibitors and SGK3 knock-out blocked phosphorylation of the expected reporters (Fig 6).

**Figure 5:**
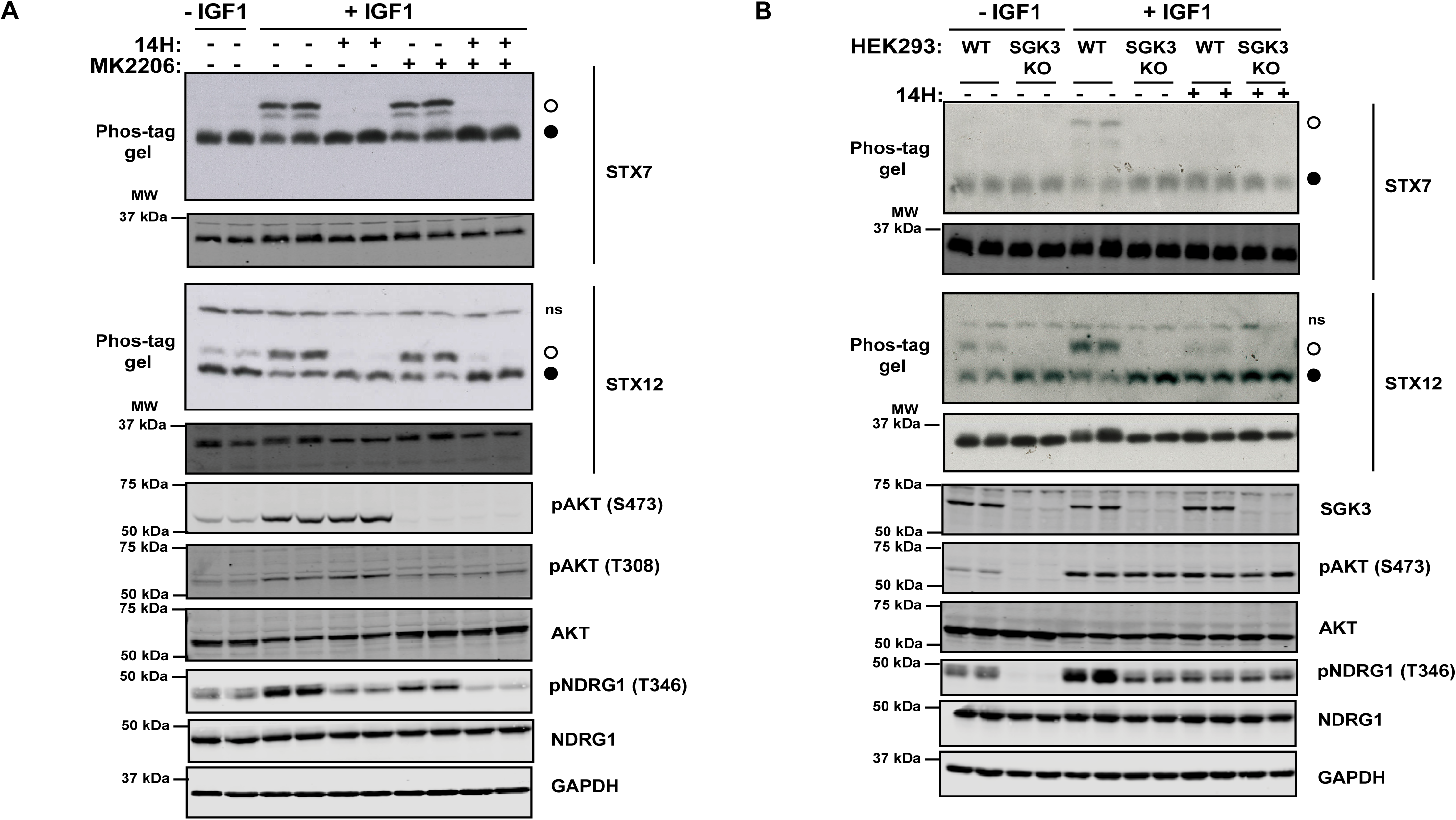
SGK3-dependent phosphorylation of endogenous STX7 and STX12 in vivo. (A) Wild type HEK293 cells were serum starved overnight, then treated for 1 h with 14H (1 µM), MK2206 (1 µM) or with DMSO, followed by stimulation with IGF1 (50 ng/ml) for 15 min. STX7 and STX12 phosphorylation was analyzed by Phos-tag assays (top panels, °□phosphorylated, ^•^□non-phosphorylated, ns; non-specific). Control immunoblots were performed on normal gels with the indicated antibodies. Immunoblots were developed using the LI-COR Odyssey CLx Western Blot imaging system analysis with the indicated antibodies at 0.5-1 µg/mL concentration.(B) Wild type (WT) and SGK3 knock-out (KO) HEK293 cells were serum starved overnight, then treated for 1 h with 14H (1 µM) or with DMSO, followed by stimulation with IGF1 (50 ng/ml) for 15 min. STX7 and STX12 phosphorylation was analyzed by Phos-tag assays (top panels, °□phosphorylated, ^•^□non-phosphorylated, ns; non-specific). Control immunoblots were performed on normal gels with the indicated antibodies.

**Figure 6:**
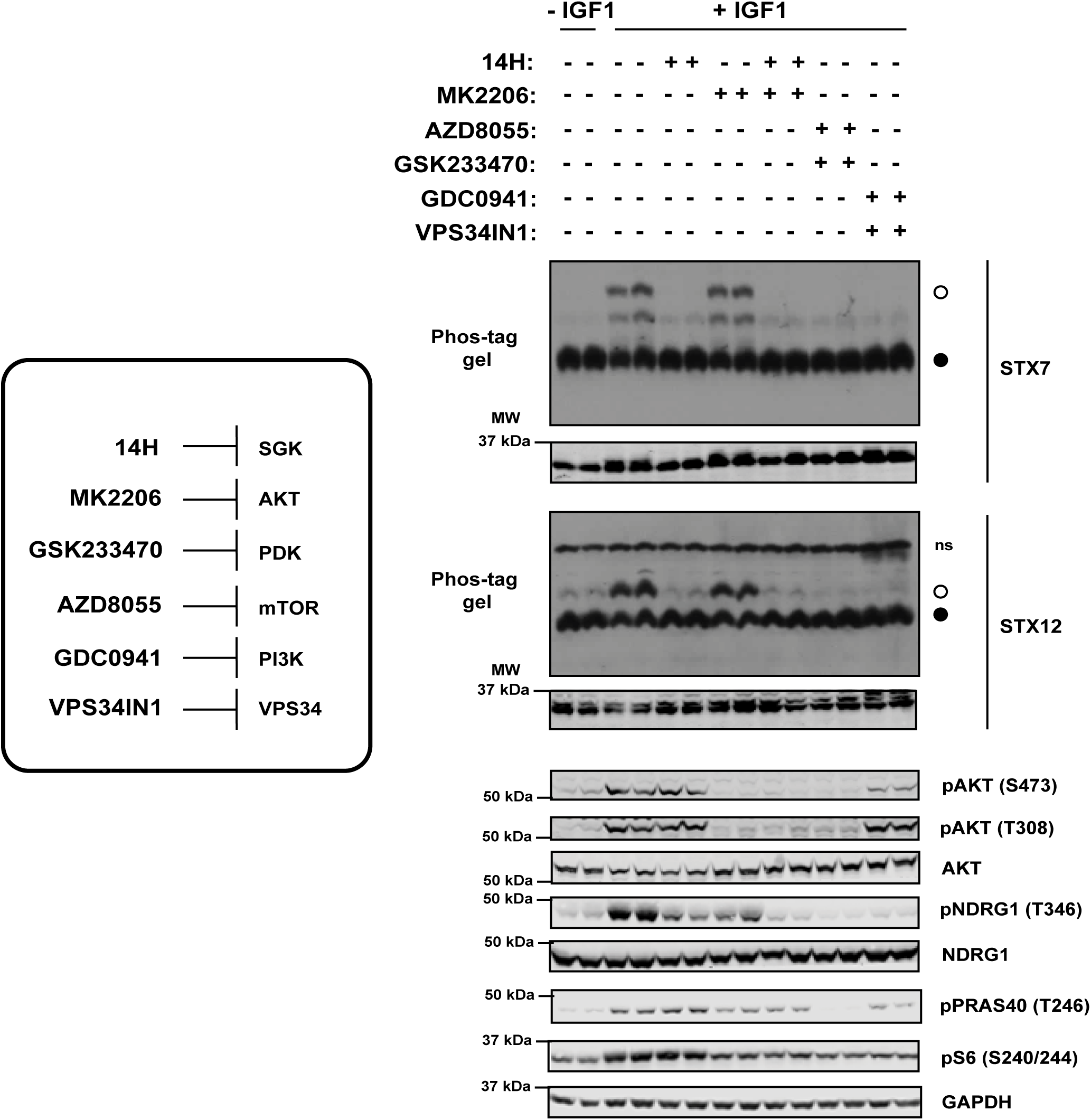
Inhibition of upstream activators of SGK3 blocks phosphorylation of STX7 and STX12 in cells. Wild type HEK293 cells were serum starved overnight, then treated for 1 h with 14H (1 µM), MK2206 (1 µM), AZD8055 (1 µM), GSK233470 (1 µM), GDC0941 (1 µM), VPS34IN1 (1 µM) or with DMSO, followed by stimulation with IGF1 (50 ng/ml) for 15 min. STX7 and STX12 phosphorylation was analyzed by Phos-tag assays (top panels, °□phosphorylated, ^•^□non-phosphorylated, ns; non-specific). Control immunoblots were performed on normal gels with the indicated antibodies. Immunoblots were developed using the LI-COR Odyssey CLx Western Blot imaging system analysis with the indicated antibodies at 0.5-1 µg/mL concentration.

Similarly, IGF1 stimulation (15 min, 50 ng/ml) also induced a major band-shift on a phos-tag gel for STX12 corresponding to ∼60% of the protein becoming phosphorylated (Fig 5A). Phosphorylation of STX12 was blocked following treatment with an SGK inhibitor (14H 1 μM) (Fig 5A) and knock-out of SGK3 (Fig 5B). Only a single shifted phosphorylated species was observed for STX12 that was also present at lower levels in serum starved cells, indicating basal phosphorylation of this site persists (Fig 5). Consistent with SGK3 mediating the phosphorylation of STX7 and STX12 treatment of cells with kinase inhibitors that suppress the activation of SGK3 namely PDK1 (GSK233470 1 µM [48]), mTOR (AZD8055 1 µM [49]), class 1 PI3K (GDC0941 1 µM [50]) and VPS34 (VPS34-IN1 1 µM [17]) blocked the IGF1 mediated band-shift of STX7 and STX12 (Fig 6). Phylogenetic analysis of the syntaxin family reveals that STX7 and STX12 are closely related and that two other syntaxin proteins possess conserved RxRXXS/T motifs, namely STX5 (Ser9) and STX16 (Ser151) (Fig S5A). However, neither of them was identified in our phosphoproteomic screens. Moreover, phos-tag analysis does not reveal detectable phosphorylation of STX5 or STX16 following IGF1 stimulation in HEK293 cells (Fig S5B).

### STX7 and STX12 are phosphorylated at Ser126 and Ser139 by SGK3 in HEK293 cells

To demonstrate that IGF1 was inducing phosphorylation of STX7 at Ser126 and STX12 Ser139, we generated STX7 and STX12 knock-out HEK293 cells employing CRISPR technology (Fig S6) and then re-expressed wild type or the respective Ala mutants of the SGK3 phosphorylation site (Fig 7). This confirmed that IGF1 failed to induce a major band-shift of the STX7[S126A] (Fig 7A) and STX12[S139A] (Fig 7B) mutants under conditions which a band-shift was observed for the wild type proteins. Phosphorylation of the minor species of STX7 was still observed in the STX7[S126A] mutant (Fig 7A), albeit slightly reduced, suggesting that the second phosphorylation site is not greatly impacted by mutation of Ser126.

**Figure 7:**
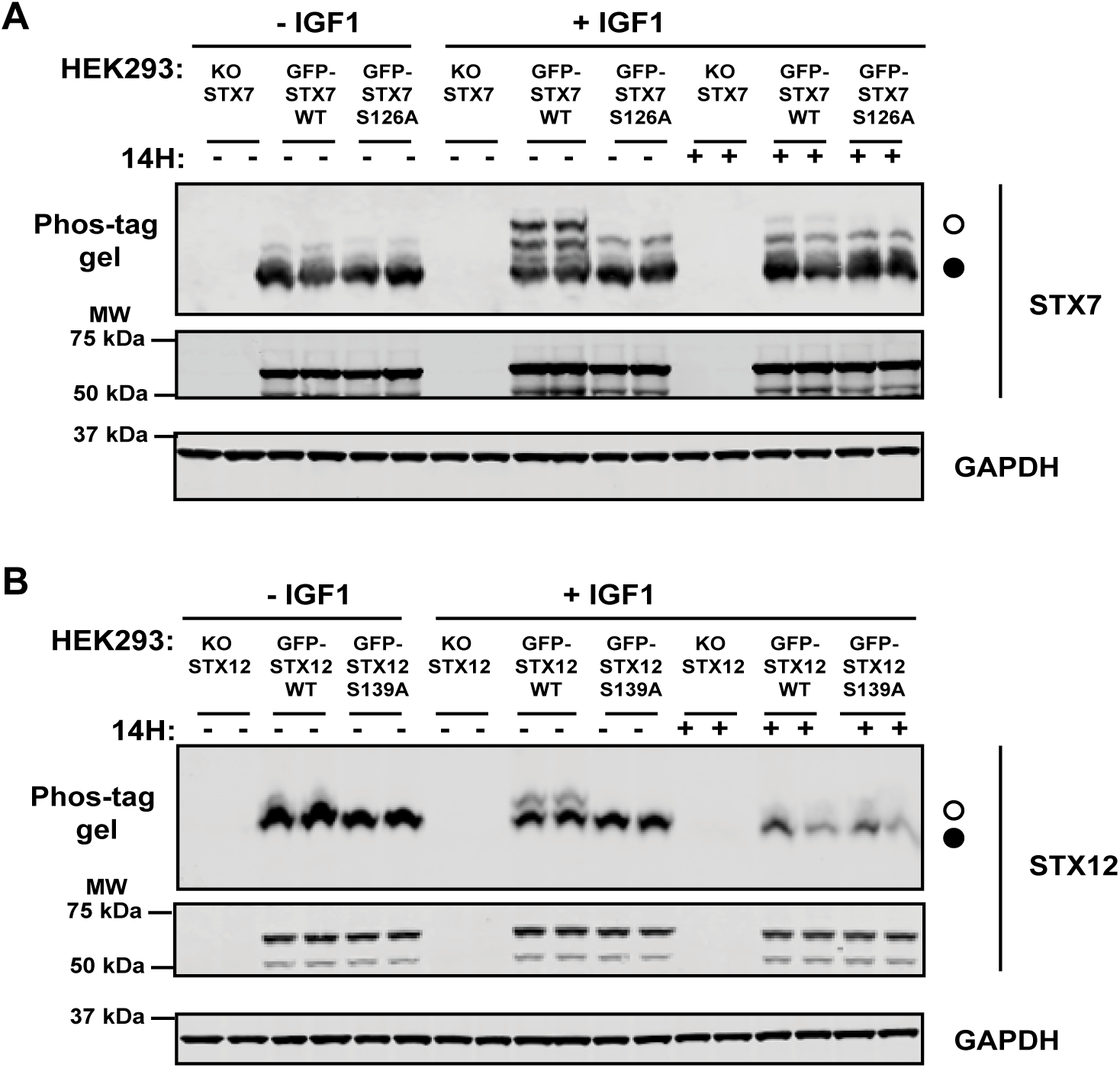
IGF-dependent phosphorylation of STX7 at Ser126 and of STX12 at Ser139 by SGK3. HEK293 stable cells lines expressing (A) GFP-STX7 wild type (WT) or the Ala mutant GFP-STX7 S126A and (B) GFP-STX12 wild type (WT) or the Ala mutant GFP-STX12 S139A were generated in STX7 knock-out (KO) and STX12 KO HEK293 cells respectively. A) KO STX7, GFP-STX7 WT and GFP-STX7 S126A HEK293 cells were serum starved overnight, treated for 1 h with 14H (1 µM) or with DMSO, followed by stimulation with IGF1 (50 ng/ml) for 15 min. STX7 phosphorylation was analyzed by Phos-tag assay (top panels, °□phosphorylated, ^•^□non-phosphorylated). Control immunoblots were performed on normal gels with the indicated antibodies. Immunoblots were developed using the LI-COR Odyssey CLx Western Blot imaging system analysis with the indicated antibodies at 0.5-1 µg/mL concentration.(B) KO STX12, GFP-STX12 WT and GFP-STX12 S139A HEK293 cells were serum starved overnight, treated for 1 h with 14H (1 µM) or with DMSO, followed by stimulation with IGF1 (50 ng/ml) for 15 min. STX12 phosphorylation was analyzed by Phos-tag assay (top panels, °□phosphorylated, ^•^□non-phosphorylated, ns; non-specific). Control immunoblots were performed on normal gels with the indicated antibodies. Immunoblots were developed using the LI-COR Odyssey CLx Western Blot imaging system analysis with the indicated antibodies at 0.5-1 µg/mL concentration.

### Evidence that Ser139 phosphorylation of STX12 enhances association with its interacting SNARE complex

Previous work has suggested that STX12 forms a complex with other SNARE proteins including STX6, VTI1a and VAMP4 and this is necessary for the fusion of early endosomes [51, 52]. To explore whether phosphorylation of STX12 at Ser139 might impact complex assembly, we immunoprecipitated GFP tagged STX12 wild type or Ser139Ala mutant that was stably expressed in STX12 knock-out HEK293 cells treated ± IGF1 (50 ng/ml, 15 min) and ± SGK inhibitor (1 µM, 1 h). The immunoprecipitated complexes were analyzed by phos-tag and conventional immunoblot analysis. IGF1 stimulation induced ∼20% phosphorylation of stably expressed wild type STX12 as judged by phos-tag analysis. STX6, VTI1a and VAMP4 were readily detected in immunoprecipitates derived from GFP-STX12 and GFP-STX12[S139A] cells but not from parental STX12 knock-out cells (that do not express GFP-STX12) (Fig 8). IGF1 stimulation reproducibly increased around 2-fold the amount of STX6, VTI1a and VAMP4 co-immunoprecipitating with wild type STX12 in three independent experiments. Mutation of the SGK3 Ser139 phosphorylation site to Ala or treatment with SGK inhibitor (14H) prevented the effect that IGF1 stimulation had on promoting association of STX12 with STX6, VTI1a and VAMP4 (Fig 8).

**Figure 8:**
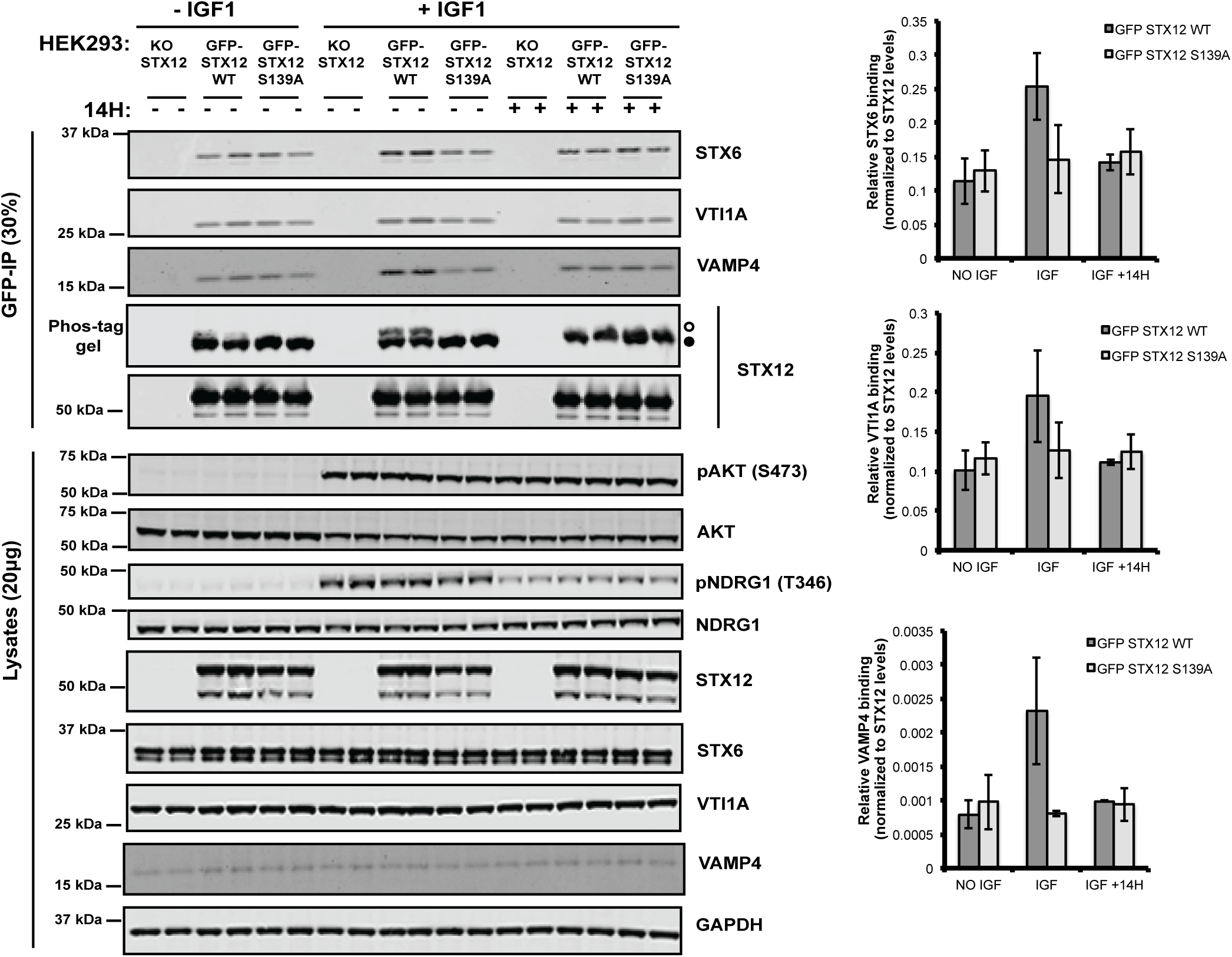
Phosphorylation of STX12 at Ser139 by SGK3 enhances the association with the SNARE complex. (A) The indicated knock-out (KO), and KO cells stably re-expressing wild type or STX[S139A] HEK293 cells were serum starved overnight, treated for 1 h with 14H (1 µM) or with DMSO, followed by stimulation with IGF1 (50 ng/ml) for 15 min. GFP-STX12 WT and GFP-STX12 S139A were immunoprecipitated using anti-GFP beads and the immunoprecipitates were subjected to western blot analysis using the indicated antibodies. STX12 phosphorylation was analyzed by Phos-tag assay (°□phosphorylated, ^•^□non-phosphorylated). Control immunoblots were performed on normal gels with the indicated antibodies. Immunoblots were developed using the LI-COR Odyssey CLx Western Blot imaging system analysis with the indicated antibodies at 0.5-1 µg/mL concentration. Quantification of the respective binding is also provided.

### Evidence that Ser139 phosphorylation of STX12 enhances localization to the plasma membrane

To investigate whether phosphorylation of STX12 by SGK3 might affect its cellular localization we performed immunofluorescence analysis of STX12 knock-out HEK293 cells that stably express either wild type GFP-STX12, the GFP-STX12 Ser139Ala mutant (Fig 9). GFP immunofluorescence of serum starved cells showed that both the wild-type STX12 and the Ser139Ala mutant were mainly localized in disperse, punctate structures that co-localize with the Rab5 early endosomal marker (Fig 9A). Moreover, in serum-starved cells expressing wild type GFP-STX12 (Fig 10B), there was a small pool STX12 present at the plasma membrane co-localizing with the Na ATPase plasma membrane marker. However, in serum starved cells expressing GFP-STX12[S139A] (Fig 9B), less plasma membrane localization of the STX12[S139A] mutant was observed. IGF1 stimulation led to a significant increase in plasma membrane localization of wild type GFP STX12 (Fig 9B), which was blocked by treatment with SGK inhibitor (14H) (Fig 9B). In contrast, IGF1 induced less plasma membrane localization of the GFP-STX12[S139A] mutant compared to the wild type respectively (Fig 9B).

**Figure 9:**
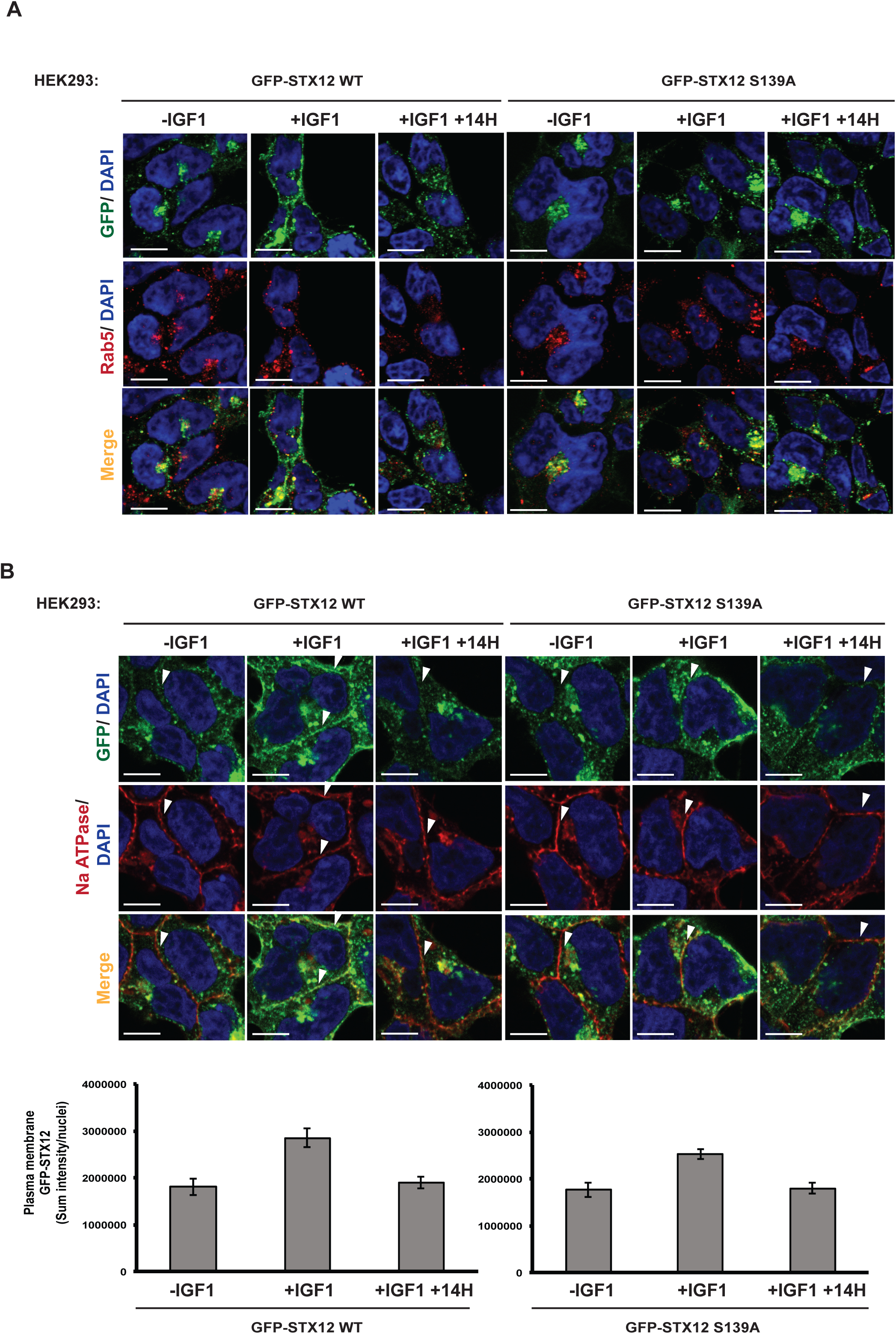
Phosphorylation of STX12 at Ser139 by SGK3 promotes its localization to the plasma membrane. STX12 knock-out (KO) HEK293 cells stably expressing GFP-STX12 WT and GFP-STX12 S139A were seeded on coverslips, serum-starved overnight and treated as indicated with or without 14H (1 µM) for 60 min prior to stimulation with IGF1 for 15 min. Cells were subsequently fixed with 4% by vol paraformaldehyde and GFP distribution was visualized using chicken anti-GFP primary and anti-chicken Alexa Fluor 488 to enhance the GFP signal. A) Co-localization of GFP tagged proteins with an early endosomal Rab5 marker was visualized using mouse anti-Rab5 primary and anti-mouse Alexa Fluor 594 secondary antibody. B) Co-localization to the plasma membrane was visualized using a rabbit anti-Na ATPase and anti-mouse Alexa Fluor 594 secondary antibody. Scale bar; 10 µm. Arrows indicate regions of Na ATPase-positive plasma membrane. Quantification of the GFP-STX12 WT or GFP-STX12 S139A plasma membrane localization is also provided.

## Discussion

To our knowledge this is the first phosphoproteomic study aimed at uncovering physiological substrates for SGK3. In the PS1 phosphoproteomic screen we identified 26 putative substrates phosphorylated at in an RxRXXS/T motif in which phosphorylation was robustly reduced in SGK3 knock-out cells. Two of these substrates were previously reported SGK isoform substrates namely NEDD4L [26-28] and NDRG1 [30]), a third substrate is NDRG3 a homologue of NDRG1 that is also likely be phosphorylated by SGK as one SGK phosphorylation site of NDRG1 is conserved in NDRG3. All ten other PS1 substrates tested, namely STX7, STX12, ITSN1, ATXN1, AFF4, BAB1, PANK4, RFIP4, MPST and WDR44, were efficiently phosphorylated *in vitro* by SGK3 at the site identified in the PS1 phosphoproteomic screen (Fig 3, S2, S3). Further work is required to investigate whether the remaining 13 PS1 substrates namely MYCBP2, CKLF4, M4K4, KIF1B, MAGI, MP2K4, NIPBL, SASH1, SRM2, ZBT32, ZN318 and ZO2 are indeed efficiently phosphorylated by SGK3.

In the PS2 phosphoproteomic screen we identified 19 potential substrates phosphorylated at in an RxRXXS/T motif in which phosphorylation was reduced following treatment with 3 μM 14H SGK isoform inhibitor, that inhibits SGK isoforms with an IC50 of 4-10 nM [12]. Although 14H is the most selective and well characterized SGK3 inhibitor, it is not possible to rule out that impact on substrate phosphorylation observed in PS2, is not a result of an off-target effect of 14H. The selectivity of 14H was assessed using a limited panel of 140 kinases and the key off target kinases were S6K1 with IC50 of 76 nM and MLK isoforms with IC50 of 100–600 nM [12]. Previous work suggested that 1 μM 14H only modestly inhibited S6K1 *in vivo* [12]. We were able to demonstrate that three out of the five substrates identified in both PS1 and PS2 were phosphorylated by SGK3 namely STX7, STX12, ITSN1 and all comprise endosomal proteins (Fig 3, S2). The fourth hit NEDD4L, was not tested, but is a well characterized SGK3 substrate [26-28]). The fifth substrate observed in both PS1 and PS2 was MYCBP2, one of the largest proteins encoded in the human proteome (4678 residues). MYCBP2 encodes an E3 ligase controlling Wallerian degeneration, namely the active process of degeneration that results when a nerve fiber is damaged and the part of the axon distal to the injury degenerates [53]. MycBP2 controls Wallerian degeneration through destabilization of nicotinamide mononucleotide adenyltransferase [54]. It has recently been shown to ubiquitylate substrates on Thr residues rather than Lys [55]. As the same MYCBP2 phosphorylation site (Ser2871) was observed in both PS1 and PS2, MYCBP2 and our data suggests it is a strong candidate for a genuine SGK substrate. Further work is required to establish this and determine whether this phosphorylation is involved in regulating E3 ligase activity and/or Wallerian degeneration. Further work is also required to assess the other 19 substrates identified in the PS2 phosphoproteomic screen that were not observed in PS1 namely AP1AR, ARFGEF1, ARFGEF2, BRSK2, CAST, CCNY, CENPA, CMTM4, IRS1, NDRG1, NHSL2, PLEC, POLR2M, WDR7. From these only NDRG1 is a known SGK isoform substrate [30] . A priority for future work would be to rule out that these phosphoryations were not due to off-target effects of 14H and confirm that SGK isoform knock-out or treatment with inhibitors that block SGK isoform activation also suppress phosphorylation of these residues. We did not identify in PS1 or PS2 phosphorylation of AIP4 [31] and FLI-1 [32] that were previously reported SGK3 selective substrates.

We focused our analysis exclusively on proteins phosphorylated within RXRXXS/T motifs that are likely to comprise direct SGK3 substrates. The mass spectrometry data set for PS1 and PS2 also contained numerous other phosphorylated substrates not lying in a RXRXXS/T motifs, whose phosphorylation is impacted by SGK3 knockout or 14H treatment. This data is available for further analysis (Supplementary Excel file) and all data from PS1 and PS2 screens have been deposited on the PRIDE database PXD014561. A motif analysis revealed potentially enriched phosphomotifs such as RXRS (7 statistically significant substrates in PS1 and 2 in PS2) or RXXS/TP (56 statistically significant substrates in PS1 and 230 in PS2) (Supplementary Excel file). Further investigation is needed to understand the mechanism by which these sites might be regulated by SGK3.

It is likely that many of the SGK3 substrates identified in PS1 and PS2 will also comprise Akt substrates. Indeed, five out of 26 substrates identified in PS1, have previously been reported as Akt substrates namely ATXN1 [39], BABA1/Merit40 [38], NDRG1 [10], NEDD4L [28] and WDR44 [40]. Furthermore, six out of the 10 substrates that were tested namely NDRG1, MST1, PANK4, ITSN1, ATXN1 and AFF4 were similarly phosphorylated *in vitro* by both Akt and SGK3. Akt has a strong requirement for a large hydrophobic residue at the n+1 position, but SGK isoforms may better tolerate diverse residues at the n+1 position [23, 33]. The majority of substrates identified in the PS1 and PS2 screens did not have a large hydrophobic residue at the n+1 position. Only five out of 26 substrates identified in PS1 had a large hydrophobic residue at the n+1 position namely KF1B-Ser1053(Leu), MAGI-Ser1443 (Leu), MP2K4-Ser80 (Ile) as well as PANK4-Thr406 (Phe) and MPST-Ser17 (Val) that we demonstrated were phosphorylated by both SGK and Akt similarly (Fig S2). Three out of 19 substrates in PS2 had a n+1 hydrophobic motif, namely ARFGEF1-Ser1079 (Leu), ARFGEF2-Ser277 (Leu) and PLEC-Ser4386 (Val) that would be predicted to be phosphorylated by both Akt and SGK. The n+1 position of the other substrates identified was Lys/Arg-9 substrates, Ser/Thr-5 substrates, Asp/Glu-4 substrates, Gly/Ala-6 substrates, Asn/Gln-4 and Pro-3 substrates (Fig 2C). Substrates with an n+1 Lys, Arg, Asp, Glu and Pro would not be predicted to be well phosphorylated by Akt [23] and are more likely to comprise SGK isoform selective substrates.

Recently, it has been reported that Akt can phosphorylate WDR44 at Ser342 and Ser344 and this phosphorylation enhances WDR44 binding to Rab11 and ultimately inhibits ciliogenesis [40]. Our data indicate that Ser344 is the phosphorylation site as mutation of Ser344 to Ala blocked phosphorylation of WDR44 (Fig 3D). We also observed that Akt phosphorylated WDR44 *in vitro* to a much lesser extent than SGK3 and it would thus be interesting to explore whether SGK3 mediated phosphorylation of WDR44 could control Rab11 and ciliogenesis . As phosphorylation of WDR44 by SGK3 and by Akt in previous work [40], has thus far only been studied *in vitro*, it would be important to generate a phospho-specific antibody that recognizes WDR44 phosphorylation at 344 to better study whether it is phosphorylated in vivo by Akt and/or SGK isoforms.

The well characterized NDRG1 substrate that is phosphorylated by both Akt and SGK possesses four phosphorylation sites that have n+1 Ser and Gly residues (Ser330 n+1=Gly, Thr346 n+1=Ser, Thr356 n+1=Ser and Thr366 n+1=Ser). It is therefore possible that substrates such as RFIP4 and WDR44 that possess a Ser or Gly at the n+1 position could indeed comprise dual Akt or SGK substrates. We have also observed that STX7 and STX12, can be phosphorylated by SGK1 as efficiently as SGK3 in vitro (Fig S4). As kinase domains of SGK kinases are so similar they are likely to have very similar intrinsic substrate specificity. The HEK293 cells that we have used for our studies express low levels of SGK1 [56] and it is possible that in cell lines that express higher levels of SGK1 or SGK2 that STX7, STX12 or other substrates identified in this study might be phosphorylated by other SGK isoforms. The four SGK3 substrates identified that were poorly phosphorylated by Akt all possessed unfavorable n+1 residues, namely STX7 (n+1 =Arg, STX12 n+1=Arg), RFIP4 (n+1=Ser) and WDR44 (n+1 = Gly). Mutation of the residue n+1 residue to Phe in all 3 of these substrates tested (STX7, STX12 and RFIP4), markedly enhanced phosphorylation by Akt without affecting SGK3 phosphorylation (Fig 4). This provides strong evidence that the n+1 reside comprises a key determinant for Akt versus SGK specificity. Other factors such as cellular co-localization at the endosome and potentially unknown docking and scaffolding interactions are likely to play important roles in mediating the selectivity of phosphorylation. STX7, STX12 and SGK3 all reside at the endosome and it us likely that also plays a major role in accounting specific phosphorylation of STX7 and STX12 by SGK3 (Fig 10). In future work it would also be interesting to explore further whether ITSN1 that is also endosomally localized is a physiological substrate for SGK3 and whether this can be also phosphorylated by Akt as suggested by our *in vitro* data (Fig S2C).

**Figure 10:**
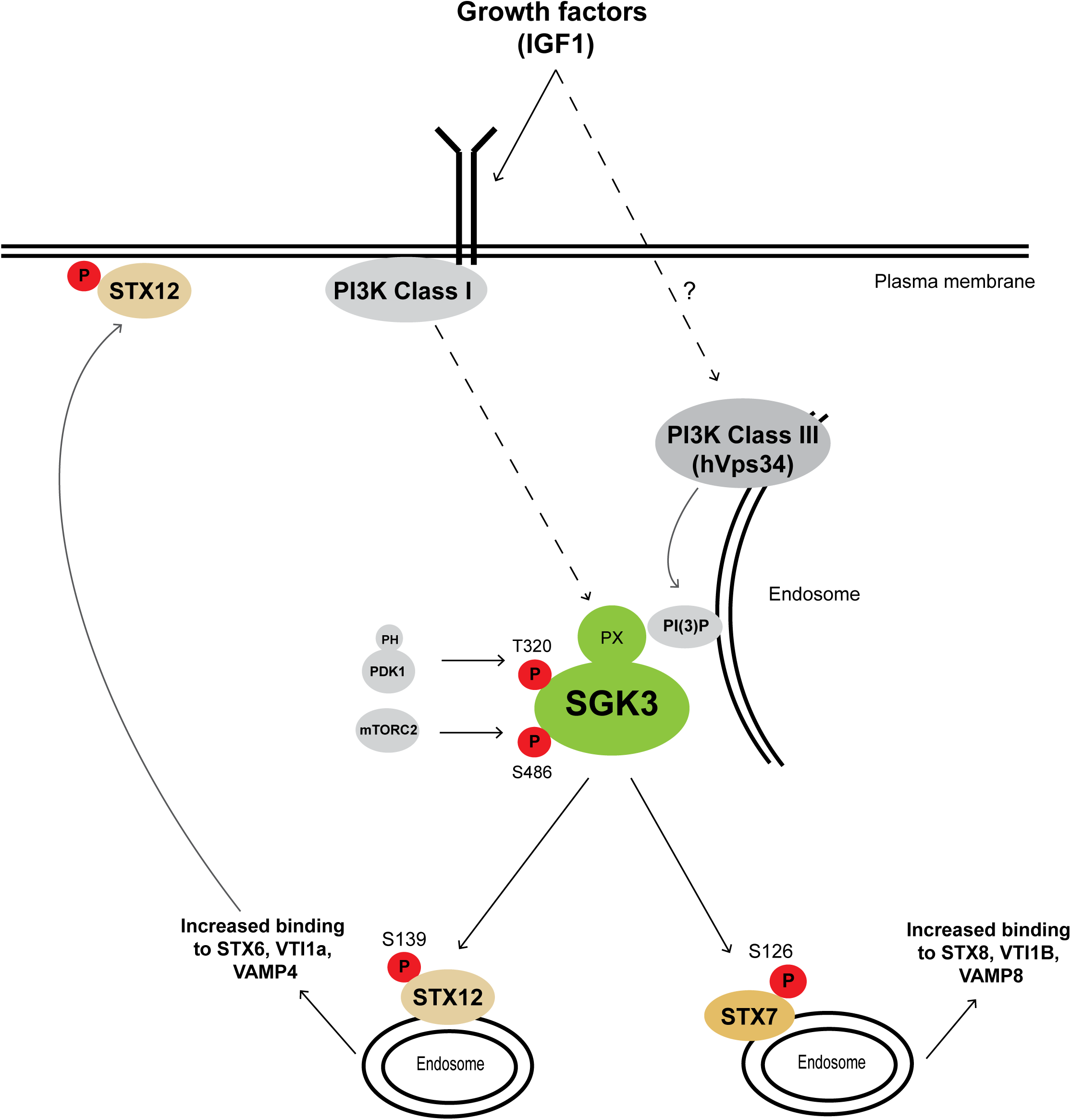
Model of the SGK3 signalling pathway. A) IGF1 stimulates production of PtdIns(3)P by the Class I and class III Vps34 at the endosomes where SGK3 can be recruited and activated by PDK1 and mTORC2 [22]. Once activated SGK3 phosphorylates its selective substrates STX7 and STX12 that are involved in the endocytic pathway. Phosphorylation of STX12 at Ser 139 by SGK3 that is located just beyond the N-terminal Habc regulatory domain promotes interaction with the t-SNARE proteins syntaxin 6 and Vti1a and the v-SNARE protein VAMP4. Our data suggest that this promotes plasma membrane localization. We propose that assessing phosphorylation of STX7 (Ser126) and STX12 (Ser139) that are specific substrates for SGK2 could be used as key biomarkers for the SGK3 signalling pathway.

Both STX7 and STX12 belong to the large family of SNARE (Soluble *N*-ethylmaleimide (NEM)-sensitive factor attachment receptor) proteins that are involved in membrane fusion in the secretory pathway. All SNARE proteins contain a highly conserved SNARE domain of about 60 amino acid residues that is responsible for the direct interaction with the SNARE domains of other members of the family [57]. There 15 members of the syntaxin family in mammals (Fig S5). They comprise an N-terminal regulatory domain termed Habc (also referred to as the STX2 domain), a SNARE domain and a C-terminal membrane-targeting sequence and with the exception of syntaxin 11, they are all transmembrane proteins. Through their SNARE domain, syntaxins can interact with other SNARE proteins on target membranes, forming a t-SNARE complex, which in turn can bind the SNARE domain of vesicle SNARE proteins (vesicle-associated membrane proteins or VAMPs) located on the cargo-vesicles and thus promoting the docking and fusion of the vesicles with the target-membranes [58]. Previous work showed that SNARE proteins are highly phosphorylated in the endo-lysosomal system [59, 60]. Phylogenetic analysis of the syntaxin family reveals that STX7 and STX12 are closely related and that two other syntaxin proteins possess conserved RxRXXS/T motifs, namely STX5 (Ser9) and STX16 (Ser151) (Fig S5). Neither of these sites on STX5 or STX16 was identified in our phosphoproteomic screen. We have undertaken phos-tag analysis of in HEK293 cells and not observed any significant phosphorylation shift of STX5 and STX16 in response to IGF1 stimulation under conditions where STX7 is well phosphorylated (Fig S5B).

STX7 is part of a well characterized SNARE complex comprising the t-SNARE proteins syntaxin 8 and Vti1b and the v-SNARE, VAMP8 and is involved in the fusion of late endosomes with lysosomes [42, 61]. Furthermore, it has been reported that CSF-1 stimulation in macrophages induced rapid serine phosphorylation of STX7 and increased its binding to STX8, Vti1b and VAMP8 [62]. It was suggested from bioinformatic and inhibitor analysis that PKC and Akt mediated this CSF-1-induced phosphorylation of Ser-125 and Ser-129, as well as the Ser126 [62], the SGK3 phosphorylation site identified in this study. Our *in vitro* and *in vivo* data suggest that SGK3 is likely to be the kinase that phosphorylates STX7 at Ser 126 as pharmacological inhibition or knocking out of SGK3 prevents STX7 phosphorylation, whereas inhibition of Akt has no effect on STX7 phosphorylation (Fig 5 to 7). Phos-tag analysis indicates that upon IGF1 stimulation STX7 gets phosphorylated at a second site that we have not mapped, which could comprise the Ser-125 and Ser-129 sites identified previously [62]. We observed that mutation of Ser126 to Ala does not impact phosphorylation of this second site (Fig 7). Further work is required to identify this residue and the upstream kinase that phosphorylates this.

To our knowledge phosphorylation of STX12 by Akt/SGK isoforms has not been previously studied. STX12 is localized in early and recycling endosomes as well as the plasma membrane, and interacts with the t-SNARE proteins syntaxin 6 and Vti1a and the v-SNARE protein VAMP4 and the formed complex is responsible for the homotypic fusion of early endosomes [51, 52]. Furthermore, it has been suggested that STX12 can function at the plasma membrane and can regulate membrane fusion and recycling of plasma membrane proteins [63]. The N-terminal Habc regulatory interacts intramolecularly with the SNARE domain, rendering the protein to a “closed conformation” and prohibiting the interaction with the cognate partners and the assembly of the SNARE complex [64-66]. Phosphorylation of various syntaxins can impact on their interaction with other SNARE-or regulatory proteins and thus on the formation of the SNARE complex [60, 67-69]. For instance, CaMKII can phosphorylate syntaxin-3, at an N-terminal site close to the Habc domain, and this phosphorylation enhances its binding to its SNARE partner, SNAP-25 [70]. Interestingly, the phosphorylation site of STX12 by SGK3 is located at the very end of the N-terminal regulatory domain Habc domain (Fig 3E). Consistent with phosphorylation enhancing SNARE complex formation, the IGF1 stimulated phosphorylation of STX12 at Ser139 by SGK3 increased the binding to the cognate SNARE partners, namely STX6, VTI1a and VAMP4, up to two-fold (Fig 8), whereas no increase was detected with the Ala mutant at this site of STX12 or after treatment with the SGK3 inhibitor 14H. Hence, we propose that SGK3 phosphorylates STX12 at Ser139 at the endosome, leading to a conformational change that renders STX12 in an open-state, facilitating thus its interaction with STX6, VTI1A and VAMP4 and the formation of the SNARE complex (Fig 10). Furthermore, our immunofluorescence data suggest that phosphorylation of STX12 by SGK3 enhances its plasma membrane localization and are in agreement with previous findings showing that STX12 functions at the plasma membrane [71, 72].

In summary, through phosphoproteomic screens we have identified the first *in vivo* SGK3 selective substrates, namely STX7 and STX12. According to our model and previous data, upon IGF1 stimulation SGK3 is activated via both PI3K Class I and III pathways and phosphorylates its selective substrates, STX7 and STX12 (Fig 10). Phosphorylation of STX7 [62] and STX12 enhances its interaction with its cognate SNARE partners and the formation of the SNARE complex. At least for STX12, this promotes recruitment to the plasma membrane (Fig 10). Moreover, we identified two other endosomal localized substrates RFIP4 and WDR44 that at least *in vitro* are preferentially phosphorylated by SGK3 and are strong candidates for comprising *in vivo* SGK3 substrates. We also identified numerous other potential new substrates for SGK3 including ITSN1 and MycBP2, that are implicated in diverse biology with links to human disease that would warrant further investigation. Our data suggest monitoring phosphorylation of STX7/STX12 or other SGK3 substrates identified in this study, could be exploited as biomarkers of SGK3 activity. Development of such biomarker assays could help identify tumors where SGK3 is a proliferation driver. Such tumors would be predicted to be resistant to Akt pathway suppressors and would instead be better treated by deploying a strategy that targets the SGK pathway.

## Materials and methods

### Materials

[γ-^32^P]ATP was purchased from PerkinElmer. Triton X-100, EGTA, sodium orthovanadate, sodium glycerophosphate, sodium fluoride, sodium pyrophosphate, 2-mercaptoethanol, sucrose, Tween 20, Triton X-100 Tris‒HCl, sodium chloride and magnesium acetate were from Sigma. PMSF was from Melford. GFP-Trap beads were purchased from Chromotek. Tissue culture reagents, Novex 4‒12% Bis‒Tris gels and NuPAGE LDS sample buffer were from Invitrogen. Polyethylenimine was from Polysciences. Polybrene was from Sigma. Methanol was from VWR Chemicals. Inhibitor GDC-0941 was from Axon Medchem and MK-2206, AZD8055 and GSK2334470 were purchased from Selleckchem. VPS34-IN1 (1-[[2-[(2-chloropyridin-4yl)amino]-4-(cyclopropylmethyl)-[4,5′-bipyrimidin]-2-yl]amino]-2-methyl-propan-2-ol) was synthesized as described in patent WO 2012085815 A1 as described before [17]. The Sanofi compound 14 h was synthesized as described recently [12]. All plasmids used in this study were generated by the MRC-PPU reagents and Services team (https://mrcppureagents.dundee.ac.uk/). All DNA constructs were verified by DNA sequencing, performed by the MRC-PPU DNA Sequencing and Service (http://www.dnaseq.co.uk). All constructs are available to request from the MRC-PPU reagents webpage (http://mrcppureagents.dundee.ac.uk), and the unique identifier (DU) numbers indicated above provide direct links to the cloning and sequence details.

### Antibodies

The following antibodies were raised in sheep, by the MRC–PPU reagents and Services team (https://mrcppureagents.dundee.ac.uk/) at the University of Dundee, and affinity purified against the indicated antigens: anti-Akt1 (S695B, third bleed; raised against residues 466–480 of human Akt1: RPHFPQFSYSASGTA), anti-NDRG1 (S276B third bleed; raised against full-length human NDRG1) (DU1557) and anti-SGK3 (S037D second bleed; raised against human SGK3 PX domain comprising residues 1–130 of SGK3) (DU2034). Anti-phospho-Akt Ser473 (#9271), anti-phospho-Akt Thr308 (#4056), anti-phospho-NDRG1 Thr346 (#5482), anti-phospho-rpS6 Ser240/244 (#2215), anti-phospho-PRAS40 Thr246 (#2997) and anti-Rab5(#46449) antibodies were purchased from Cell Signalling Technology. Anti-STX6 (10841-1-AP), anti-STX7 (12322-1-AP), anti-STX12 (14259-1-AP), anti-STX16 (11201-1-AP), anti-VAMP4 (10738-1-AP) andanti-VTI1A (12354-1-AP) were purchased from Proteintech. Anti-STX5 (ab217130), anti-GFP(ab13970) and anti-Na ATPase (ab76020) were purchased from Abcam. Anti-GAPDH (sc-32233) was from Santa Cruz Biotechnology. Secondary antibodies coupled to IRDye680LT or IRDye800CW were obtained from Licor Biosciences. Secondary antibodies coupled to HRP (horseradish peroxidase) were purchased from Thermo Scientific. Secondary antibodies coupled to Alexa Fluor dyes were purchased from Life technologies (Invitrogen).

### Cell culture and cell lysis

HEK293, cells were purchased from the American Tissue Culture Collection (ATCC). Cells were cultured DMEM media supplemented with 10% by vol FBS, 2 mM L-glutamine, 100 U/ml penicillin and 0.1 mg/ml streptomycin. Inhibitor treatments were performed as described in figure legends The cells were lysed on ice in lysis buffer containing 50 mM Tris–HCl (pH 7.5), 1 mM EGTA, 1 mM sodium orthovanadate, 10 mM sodium glycerophosphate, 10 mM sodium pyrophosphate, 50 mM sodium fluoride, 0.27 M sucrose, 1 mM DTT, 1 mM benzamidine and 0.1 mM PMSF. Lysates were clarified by centrifugation at 11,000 *g* for 20 min at 4°C. Protein concentration was estimated by Bradford assay (Thermo Scientific). Immunoblotting and immunoprecipitation were performed using standard procedures. The signal was detected using a Li-Cor Biosciences Odyssey System and quantified in Image Studio Lite (Li-Cor) or using ECL Western Blotting Detection Kit (Amersham) on Amersham Hyperfilm ECL films (Amersham).

### Phosphopeptide enrichment and Tandem Mass tags labelling

For PS1, SGK3 knock out HEK 293 (SGK3 knock-out) cells were generated by Crispr/Cas9 methodology as described earlier. Wild type and SGK3 knock-out cells were treated as described in figure legends and lysed using a 2% SDS lysis buffer (2% by mass SDS, 250 mM NaCl, 50 mM HEPES pH 8.5, 1 mM Benzamidine, 2 mM PMSF, 2 mM Sodium orthovanadate, 10 mM sodium β-glycerophosphate, 5 mM sodium pyrophosphate, 50 mM sodium fluoride, supplemented with protease inhibitor cocktail tablets (Roche) and PhosSTOP phosphatase inhibitors (Roche). Lysates were heated at 95°C for 5 min prior to sonication and clarification at 14, 000 rpm for 15 min. Following determination of protein concentration by BCA assay, 25 mg protein was subjected to acetone precipitation. The extracted pellet was resolubilized in 6 M urea/50 mM triethylammonium bicarbonate (TEAB) by sonication and protein concentration determined again by BCA assay. Protein samples were subsequently reduced with 10 mM DTT and incubated at 56°C for 20 min. Following cooling, samples were alkylated with 30 mM iodoacetamide for 30 min in the dark at room temperature prior to reducing the samples again with 5 mM DTT for 10 minutes at room temperature. Protein lysates were diluted to 1.5 M urea and digested with LysC (Wako, Japan) in a 1:200 enzyme:protein ratio overnight at room temperature. Protein extracts were diluted further to a 0.75 M urea concentration, and trypsin (Promega, WI USA) was added to a final 1:200 enzyme:protein ratio for 16 h at 37 °C. Digests were acidified by the addition of trifluoroacetic acid to a final concentration of 1% by vol trifluoroacetic acid. Samples were centrifuged at 4000 RPM for 15 min at 4°C and the undigested precipitate and excess trypsin were discarded, while the supernatant retained. Samples were subsequently subjected to C18 solid-phase extraction (SPE) (Sep-Pak, Waters, Milford, MA) to remove salts and impurities. Briefly Sep-Pak cartridges were activated by adding 4 ml 100% Acetonitrile and equilibrated using 0.1% by vol trifluoroacetic acid by (2 x 4 ml). The acidified peptide digest was loaded on to the C18 cartridges. Peptides were cleaned with 2 x 4 ml 0.1% by vol trifluoroacetic acid. Peptides were subsequently eluted with 0.5 ml 60% by vol acetonitrile in 0.1% by vol trifluoroacetic acid. Finally, eluted peptides were lyophilized.

For PS2, HEK293 cells were treated with DMSO, 14H and MK2206 as described in figure legends. The cells were lysed in the same lysis buffer that was used in PS1, and 10 mg of protein amount was prepared for the Lys-C and trypsin digestion as described above and desalted the peptides same as above. 5% of the eluate was aliquoted for total proteomic analysis in both PS1 and PS2.

### Phosphopeptide enrichment

For phosphopeptide enrichment, titanium oxide (TiO_2_) beads (Titansphere Phos-TiO_2_ Bulk 10 μm #5010-21315, GL Sciences, Japan) were used [59, 73] and prepared by washing with 100% acetonitrile. Tryptic peptides were resuspended in 2 M lactic acid/50% by vol acetonitrile (pH 1.5) by water-bath sonication and centrifuged at 14,000 rpm for 15 minutes at room temperature, leaving a small pellet consisting of salts and impurities. The supernatant was transferred to fresh Eppendorf tubes and TiO_2_ beads added at a 1:2 peptide:bead ratio (For PS1, ∼20 mg of peptide digest with 40 mg of TiO_2_ beads and for PS2, ∼8 mg of peptide digest with 16 mg of TiO_2_ bead was used). The bead/peptide mixture was incubated on a mixer at 2000 rpm for 1 h at room temperature followed by centrifugation. Beads were washed twice with 500 μl of 2 M lactic acid/50% by vol acetonitrile (pH 1.5) and once with 500 μl of 0.1% by vol trufluoroacetic acid in 50% by vol acetonitrile (pH 2-4). The beads were resuspended in 200 μl of 0.1% by vol trifluoroacetic acid and loaded onto C8 stage-tips (3M Empore, #2214, USA) to separate the beads from the flow through which was discarded. Phosphopeptides were eluted from beads with 30 μl of elution buffer (50 mM potassium dihydrogen phosphate (K_2_HPO_4_)/5% by mass NH_4_OH) into Eppendorf tubes containing 6% by vol trifluoroacetic acid to quench the ammonia solution. The elution step was repeated another two times and the eluate was then subjected to C18 SepPak stage-tip purification. Briefly, four C18 disks (PN: 2215, 3M Empore, USA) were packed with 18-gauge needle into a 200 μl pipette tips. The Stage-tips were activated with 100 μl of 100% acetonitrile and then equilibrated twice by adding 100 ul of Solvent-A (0.1% by vol trifluoroacetic acid). Acidified Phosphopeptides (100 μl) were loaded twice and washed twice with 100 μl of Solvent-A and then the phosphopeptides were eluted by adding 60 μl of solvent-B (50% by vol acetonitrile in 0.1% by vol trifluoroacetic acid). The elution was repeated again the eluates were vacuum dried and continued with TMT labeling.

### Tandem Mass Tag (TMT) labelling

To carry out TMT labelling for both PS1 and PS2 as depicted in Fig 1, the TMT reagents (Thermo Fisher Scientific, TMT 10 plex kit, 0.8 mg Part number:90110) was dissolved in 100 μl anhydrous acetonitrile; phosphopeptide samples were dissolved in 140 μl of 100 mM TEAB buffer. 30 μl anhydrous acetonitrile and 30 μl of the appropriate TMT label (dissolved in acetonitrile) was added to the phosphopeptide samples (140 μl) such that the final acetonitrile concentration was 30% by vol (total volume = 200 μl). For the total proteome, 20 μl anhydrous acetonitrile and 40 μl TMT reagent (dissolved in acetonitrile) was added to 140 μl peptide sample. The samples were incubated for 2 hours at room temperature and vortexed every 30 min after which 10% was aliquoted from each sample to check the labeling efficiency on Orbitrap Fusion Tribrid mass spectrometer. Further, the reactions were quenched with 5 μl 5% by vol hydroxylamine and experimental samples were combined in equal and the multiplexed peptide and phosphopeptide samples were lyophilized and subjected to C18 Stage-tip desalting as described above.

### Basic pH Reverse-phased Fractionation

In the current study, both the TMT labelled Phosphoproteome and total proteome was subjected to basic-pH reversed phase liquid chromatography (bRPLC) fractionation. Labelled phosphopeptides were solubilized in buffer A (10 mM ammonium formate, pH 10) and separated on an XBridge BEH RPLC column (Waters XBridge BEH C18 Column, 130., 5 μm, 4.6 mm X 250 mm. #186003010) at a flow rate of 0.3 ml/min by applying a non-linear gradient of 7% to 40% (Solvent B, 90% by vol acetonitrile in 10 mM ammonium formate, pH 10) for 80 minutes into a total of 96 fractions. The 96 fractions were concatenated into 13 fractions for PS1 and 12 fractions for PS2 in a checkerboard manner and acidified with 30 μL of 1% by vol formic acid. The fractions were subsequently vacuum-dried and stored at −80°C until the LC-MS/MS analysis. The total proteome samples from both PS1 and PS2 were also fractionated and subsequently vacuum dried and stored at −80°C deep freezer until the LC-MS/MS analysis.

### LC-MS/MS analysis

Both the PS1 and PS2 phopshoproteome and total proteome samples are analyzed on Orbitrap Fusion Tribrid Mass spectrometer (Thermo Fisher Scientific, San Jose, USA) that was in line with the Dionex 3000 RSLC nano-liquid chromatography system, similar to previously published [74-76]. The peptide samples were loaded on a nano viper trap column (C18, 5u, 100Å, 100um X 2 cm; PN: 164562, Thermo Scientific) and resolved on a 50 cm analytical column (2 um, 100 Å, 75 um X 50 cm. PN: ES803, Thermo Scientific) and directly electrosprayed using Easy nano LC source (Thermo Fisher Scientific, San Jose USA) into the mass spectrometer. Both the TMT labeled phosphoproteome and total proteome fractions from PS1 and PS2 (Phosphoproteome and total proteome) were injected as technical replicates. Phosphopeptide fractions were dissolved in 15 μl of Solvent-A buffer (0.1% by vol Formic acid (FA) in 3% by vol acetonitrile). The Dionex autosampler loaded the peptides into the trap column at 10 μl/min flow rate using Solvent-A (0.1% by vol formic acid) following the peptide samples were resolved on an analytical column by applying a 3% to 25% of solvent-B at 300 nl/min flow rate (Solvent-B: 80% by vol acetonitrile in 0.1% by vol formic acid and 3% by vol DMSO) for 180 min and 25% to 35% solvent-B for 30 min and 35% to 99% Solvent-B for 5 min which was then maintained at 99% Solvent-B for 10 min and washed with Solvent-A for another 15 min comprising a total run time of 240 min. The mass spectrometer was operated in data dependent mode in a top speed comprising 2.5 sec cycle time (MS1 and MS2). The MS1 data was acquired in a scan range of 400-1600 m/z at 120,000 resolution at 200 m/z and measured using ultra high-field Orbitrap mass analyzer. The precursor ions were isolated using quadrupole with an isolation window of 1.6 m/z which are then fragmented using 37.5% beam-type collisional induced dissociation (HCD) and acquired at 60,000 resolution at 200 m/z using ultra high-filed Orbitrap mass analyzer. The AGC and ion injection times for MS1: 3E5 and 50 ms and for MS2: 5E4 and 250 ms were used. The dynamic exclusion was enabled for 35 sec to avoid the repeated sampling of the precursor ions.

### Mass spectrometry data analysis

The mass spectrometry raw data was processed for the protein identification and TMT-based quantification using MaxQuant pipeline [77, 78]. The PS1 total proteome and phosphoproteome was searched using MaxQuant version 1.5.1.7 against Human Uniprot database (release 2017-02; 42,101 sequences). The default common contaminants were selected within the MaxQuant suite. Group specific parameters: Reporter ion MS2, trypsin as a protease with a maximum of 2 missed cleavages were allowed. Acetyl Protein N-term, Deamidation of NQ, Oxidation of M and Phosphorylation of STY were selected as variable modifications (Phospho STY was excluded for total proteome). Carbamidomethylation of Cys was selected as fixed modification. First search tolerance of 20 ppm and main search tolerance of 5 ppm are allowed. Global parameters: Minimum peptide length of 7 amino acids and minimum score of 40 and minimum delta score of 6 for modified peptides was selected. 1% FDR at PSM, peptide and protein level was applied to generate the final output tables. Modified and corresponding unmodified peptides for phospho sites were excluded for accurate protein level quantification. For PS2 both the total proteome and phosphoproteome was processed using MaxQuant version 1.6.0.13 and searched against Human Uniprot database (release 2017-02; 42,101 sequences). The default common contaminants were selected within the MaxQuant suite. Two group specific parameters were selected. Group 0 for Phosphoproteome and Group 1 for total proteome. Group 0 parameters: Reporter ion MS2 with a minimum reporter PIF (Precursor intensity fraction) value of 0.75 was selected to filter out the contaminating peptide fraction for TMT quantification. Acetyl Protein N-term, Deamidation of NQ, Oxidation of M and Phosphorylation of STY were selected as variable modifications. Carbamidomethylation of Cys was selected as fixed modification. First search tolerance of 20 ppm and main search tolerance of 5 ppm are allowed. For Group 1, the phosphorylation of STY as a variable modification was excluded. Global parameters: Minimum peptide length of 7 amino acids and minimum score of 40 and minimum delta score of 6 for modified peptides was selected. 1% FDR at PSM, peptide and protein level was applied to generate the final output tables. Modified and corresponding unmodified peptides for phospho sites were excluded for accurate protein level quantification. The MaxQuant output tables, Phospho STY sites table and the protein tables were processed separately using Perseus software [79] suite. Common contaminants and reverse hits were filtered. The TMT quant intensities were log2 transformed. The TMT triplicates channels were categorized based on the sample conditions (PS1: wild type and knock-out) and (PS2: DMSO, 1 μM MK2206 and 3 μM 14H). Unquantified proteins were filtered Two tailed independent T-test by applying 5% permutation-based FDR correction was applied to identify the differentially regulated and statistically significant phosphosites and proteins. Phosphosites with P <0.05 and a fold change of 1.5-fold were considered as significantly altered in between the samples.

### Data availability

The mass spectrometry raw data and MaxQuant output tables from both the screens can be accessed using the PRIDE repository identifier PXD014561. (https://www.ebi.ac.uk/pride/archive/). Reviewer login details: Username. Password: bwyJsZSf

### Generation of CRISPR-CAS9 knock-out cell lines

The SGK3 HEK293 knock-out cell line has been described previously [22]. To generate STX7 and STX12 HEK293 knockout cell lines a modified Cas9 nickase system and the following constructs were used (available at https://mrcppureagents.dundee.ac.uk): DU60563/DU60564 (STX7), DU60130/DU60132 (STX12). The sgRNA pairs were identified with a low combined off-targeting score [STX7 knock-out-sgRNA1: GACCCCGCCCAGTTGGCCCAG (DU60563); sgRNA2: GAACTCCTGGAGTGTAAGACA (DU60564), STX12 knock-out-sgRNA1: GCCCTCGGGGCCCCAGCTCC (DU60130); sgRNA2: GGGTTCCGGTACATGTCTAA (DU60132)] using the Sanger Centre CRISPR finder tool (http://www.sanger.ac.uk/htgt/wge/find_crisprs). For STX7 knock-out the guide RNAs target exon 2 of *STX7* gene, while for STX12 knock-out the guide RNAs target exon 1. The antisense guides (sgRNA2) were cloned onto the spCas9 D10A-expressing pX335 vector (Addgene plasmid no. 42335) and the sense guides (sgRNA1) into the puromycin-selectable pBABED P U6 plasmid (Dundee-modified version of the original Cell Biolabs pBABE plasmid, DU48788). Cells at about 80% confluency were co-transfected in 10-cm dishes with 2 µg each of the sgRNA plasmid pairs and 20 µl of PEI stock solution (1 mg/ml). 24 h post transfection, the medium was replaced with fresh containing 2 µg/ml puromycin in order to select for the cells that have been transfected. After 48 h of selection, the medium was replaced with normal medium and the cells were left to recover for 24 h before performing single-cell sorting using an Influx cell sorter (Becton Dickinson). Single cells were inoculated in individual wells of a 96-well plate containing DMEM supplemented with 20% by vol FBS, 2 mM L-glutamine, 100 units/ml penicillin and 100 mg/ml streptomycin. At about 80% confluency individual clones were transferred into six-well plates and screened for STX7 or STX12 knock-out by western blotting.

### Generation of GFP-STX7 WT, GFP-STX7 S126A, GFP-STX12 WT, GFP-STX12 S139A HEK293 stable cell lines

To generate stable HEK293 cell lines requiring lentiviral infection, the following pLVX Hygro GFP constructs carrying the gene of interest were used: DU62665 (GFP-STX7 WT), DU62666 (GFP-STX6 S126A), DU62667 (GFP-STX12 WT) and DU62668 (GFP-STX12 S139A). To generate lentiviral particles, 2 μg parent plasmid (containing the gene of interest), 1.5 μg psPAX2 packaging plasmid and 500 ng pMD2.G envelope plasmid were co-transfected in HEK293 FT cells in 10-cm dishes using Lipofectamine (Invitrogen) according to manufacturer’s instructions. After 12 h of transfection, the media was replaced with fresh and the cells were cultured for 24 h. After 24 h, the media containing the lentiviral particles were collected and filtered through a 0.45 μm filter. The virus was either frozen and stored at −80°C for later use or immediately used to infect cell cultures. 1-5 ml of virus was applied directly to either HEK293 knock-out STX7 or HEK293 knock-out STX12 cells in the presence of 10 μg/ml polybrene (Hesse et al., 1978). 24 h post infection, the medium was replaced with fresh and after 24 h it was changed to selection medium containing 100 μg/ml Hygromycin B in order to select for the transduced cells.

### *In vitro* kinase assays

1 μg of kinase was incubated with 0.5-2 μg of purified protein substrate in the presence of 0.1 mM [γ^32^P]-ATP and 10 mM magnesium acetate in Buffer A (50 mM Tris-Cl pH7.5, 0.1 mM EGTA) at 30°C with shaking at 1100r.p.m for 30 min. Reactions were terminated with LDS sample buffer and resolved by SDS-PAGE electrophoresis. Proteins were detected with Coomassie staining. The gels were imaged with an EPSON scanner, then sandwiched and secured between two sheets of pre-wet cellophane (Bio-Rad) and subsequently dried in a GelAir dryer for 45-60 min. Dried gels were exposed to Amersham Hyperfilm MP overnight, in an autoradiography cassette and the films were later developed using a Konica auto-developer.

### Phosphosite identification by MS and Edman sequencing

Bacterially purified substrate proteins (1 μg) were phosphorylated using recombinant kinase (0.5 μg) in a buffer containing 50 mM Tris-HCl pH 7.5, 0.1 mM EGTA, 10 mM MgAc, 0.1 mM [γ-^32^P]ATP for 80 min at 30°C per reaction. 5-10 reactions were carried out per phospho-mapping experiment. Reactions were stopped with LDS sample buffer resolved by SDS-PAGE electrophoresis. Gels were stained with Coomassie blue to stain protein bands. The bands corresponding to the substrate protein were excised and subsequently, washed first with water, then shrunk with acetonitrile and re-swell with 50 mM Tris HCl pH 8.0. Afterwards, samples were reduced with 5 mM DTT in 50 mM Tris HCl pH 8.0 at 65°C for 20 minutes and then alkylated with 20 mM iodoacetamide in 50 mM Tris HCl pH 8.0. The gel pieces were then washed for 10 min first in water, then in 50 mM ammonium bicarbonate and finally in 50 mM ammonium bicarbonate 50% by vol acetonitrile. The gel pieces were shrunk in acetonitrile, re-swollen in 50 mM triethylammonium bicarbonate and then re-shrunk in acetonitrile. Samples were digested overnight with trypsin (5 μg/ml in 50 mM Triethylammonium bicarbonate) at 30°C and the peptides were separated on a reverse-phase HPLC Vydac C18 column (Separations Group) with an on-line radioactivity detector. The column was equilibrated in 0.1% by vol trifluoroacetic acid and developed with a linear acetonitrile gradient at a flow rate of 0.2 ml/min. Fractions (0.1 ml each) were collected and analyzed for ^32^P radioactivity by Cerenkov counting with a tricarb scintillation counter. Isolated phosphopeptide fractions were analyzed by liquid chromatography (LC)-MS/MS using a Thermo U3000 RSLC nano liquid chromatography system (Thermo Fisher Scientific) coupled to a Thermo LTQ-Orbitrap Velos mass spectrometer (Thermo Fisher Scientific) to determine the primary sequence of the phosphopeptides. Data files were searched using Mascot run on an in-house system against a database containing the appropriate substrate sequences, with a 10 ppm mass accuracy for precursor ions, a 0.6 Da tolerance for fragment ions, and allowing for Phospho (ST), Phospho (Y), Oxidation (M), and Dioxidation (M) as variable modifications. Individual MS/MS spectra were inspected using Xcalibur 2.2 (Thermo FisherScientific), and Proteome Discoverer with phosphoRS 3.1 (Thermo Fisher Scientific) was used to assist with phosphosite assignment. The site of phosphorylation of 32P-labeled peptides was determined by solid-phase Edman degradation on a Shimadzu PPSQ33A Sequencer of the peptide coupled to 80 Sequelon-AA membrane (Applied Biosystems).

### Phos-tag SDS/PAGE and immunoblot analysis

Cell lysates were mixed with 4× SDS/PAGE sample buffer [250 mM Tris/HCl, pH 6.8, 8% by mass SDS, 40% by vol glycerol, 0.02% by mass Bromophenol Blue and 4% by vol 2-mercaptoethanol] and heated at 95°C for 5 min. Samples were supplemented with 10 mM MnCl_2_ before boiling and then centrifuged for 1 min at 14,000 r.p.m. Phos-tag SDS/PAGE was carried out as described previously [<u>45</u>] with some modifications. Gels for Phos-tag SDS/PAGE consisted of a stacking gel [4% by mass acrylamide, 125 mM Tris/HCl, pH 6.8, 0.1% by mass SDS, 0.2% by vol *N,N,N*′,*N*′_tetramethylethylenediamine (TEMED) and 0.08% by mass ammonium persulfate (APS)] and a separating gel [8% by mass acrylamide, 375 mM Tris/HCl, pH 8.8, 0.1% (by mass SDS, 100 μM Phos-tag acrylamide, 200 μM MnCl_2_, 0.1% by vol TEMED and 0.05% by mass APS]. 10–40 μg samples were loaded and electrophoresed at 70 V for the stacking part (approximately 2 h) and then 120 V for 150 min with the running buffer [25 mM Tris/HCl, 192 mM glycine and 0.1% by mass SDS]. For immunoblot analysis, gels were washed for 10 min in the transfer buffer [48 mM Tris/HCl, 39 mM glycine and 20% by vol methanol] containing 10 mM EDTA and 0.05% my mass SDS three times, followed by one wash in the transfer buffer containing 0.05% SDS for 10 min. Proteins were electrophoretically transferred onto nitrocellulose membranes (Amersham Protran 0.45 μm NC; GE Healthcare) at 100 V for 180 min on ice in the transfer buffer without SDS/EDTA. Transferred membranes were blocked with 5% by mass non-fat dry milk dissolved in TBS-T [20 mM Tris/HCl, pH 7.5, 150 mM NaCl and 0.1% by vol Tween 20] at room temperature for 45 min. Membranes were then incubated with primary antibodies overnight at 4°C. After washing membranes in TBS-T, membranes were incubated with horseradish peroxidase-labelled secondary antibodies at room temperature for 1 h. After washing membranes in TBS-T, protein bands were detected by exposing Amersham Hyperfilm to the membranes using an ECL solution Amersham ECL Western Blotting Detection Reagents (GE Healthcare).

### Co-Immunoprecipitation

Cells were lysed in standard lysis buffer as described above. GFP-STX7 WT, GFP-STX7 S126A, GFP-STX12 WT or GFP-STX12 S139A, were immunoprecipitated from 2 mg of lysates using GFP-Trap beads (Chromotek) at 4°C for 4h under rotation. Subsequently, the beads were washed twice with lysis buffer containing 0.45 M NaCl for 2 min at 4 °C under rotation and then once with lysis buffer containing 0.15 M NaCl. Protein complexes were eluted from beads using 2X LDS Sample Buffer with 2% by vol beta-mercaptoethanol, and the co-immunoprecipitating proteins were detected by immunoblot analysis.

### Immunofluorescence

Wild type and knock-out STX12 HEK293 cells that stably express wild type GFP-STX12, GFP-STX12 S139A or GFP were seeded on coverslips (precoated with poly-L-Lysine) and after 48h they were serum-starved overnight. Following treatments described in figure legends, cells were fixed with 4% by vol paraformaldehyde and permeabilized with 1% by vol NP-40. Cells were blocked using 1% bovine serum albumin (BSA) in phosphate-buffered-saline (PBS), then incubated for 1 h with primary antibodies, washed three times in 0.2%by mass BSA in PBS, and incubated for 1 h with secondary antibodies. For visualization of GFP-STX12 WT and GFP-STX12 S139A the GFP signal was enhanced using a chicken anti-GFP antibody (Abcam) followed by anti-chicken secondary antibody conjugated to Alexa Fluor 488. For localization of endosomal compartments, Rab5 was stained with anti-Rab5 antibody (CST) and secondary anti-mouse conjugated to Alexa Fluor 594. For localization to plasma membrane, an anti-Na ATPase (Abcam) and anti-rabbit conjugated to Alexa Fluor 594 were used. After incubation with the secondary antibodies, the coverslips were washed three times with 0.2% BSA in PBS. Coverslips were washed once more in water and mounted on slides using ProLong Gold Antifade (ThermoFisher). The images were collected on an LSM710 laser scanning confocal microscope (Carl Zeiss) using the ×63 Plan-Apochromat objective (NA 1.4), using a pinhole chosen to provide a uniform 0.8 um optical section thickness in all the fluorescence channels.

<u>Identification of plasma membrane positive structures in cells was performed with Volocity</u> <u>(Quorum Technologies). In brief, the</u> Na ATPase positive structures (Alexa 594) were found using the Find objects method (Objects >2.5 SD from mean image intensity) while the nuclei were found using the Automatic method (Otsu’s method), nuclei were separated by using the separate touching objects step (object size 80 um^2)^ and small structures were excluded (<30um^2^). The summed GFP (488 signal) pixel intensities for the plasma membrane objects were normalised to the number of cells in the image by dividing by the number of nuclei in the image. Error bars show SEM for images (10 images per treatment).

### Statistical analysis

All experiments were performed were performed at least in duplicates and at least twice. Each figure legend describes the statistical analysis that was performed in more detail.

## Acknowledgements

We acknowledge the excellent technical support of the MRC-Protein Phosphorylation and Ubiquitylation Unit (PPU) DNA Sequencing Service (coordinated by Gary Hunter), the cloning team (coordinated by Rachel Toth), the PPU tissue culture team (coordinated by E. Allen), the Reagents and Services antibody purification teams (coordinated by H. McLauchlan and J. Hastie), the Flow Cytometry facility (coordinated by Rosemary Clarke) and the Dundee Imaging Facility. This work was supported by the Medical Research Council [grant number MC_UU_12016/2 (to D.R.A.) and MC_UU_12016/5 (to MT)] and the pharmaceutical companies supporting the Division of Signal Transduction Therapy Unit (Boehringer-Ingelheim, GlaxoSmithKline, and Merck KGaA, to D.R.A.).

## Author contribution

NM designed, executed experiments (Figures 1, 2, 3B, 3C, 3D, S1, S2A, S2B, S2D, S2E, 10) analyzed data and played a major role in interpretation of data; RSN designed, executed experiments and oversaw experiments and analysis of data performed in PS2 (Figures 1, 2, S1) analyzed data and played major role in interpretation of data; JP designed, executed experiments (Figures 1, 2) and analyzed data; TM designed and generated all guides for the CRISPR/CAS9 studies; MW undertook most of the cloning studies; ARP took and analyzed images for Figure 9 RB executed experiment Figure S3; MT directed the design of experiments and analysis of data performed in PS1 (Figures 1, 2); AK designed, executed experiments (Figures 3A, 3E, 3F, 4, 5, 6, 7, 8, 9, 10, S2C, S2F, S4, S5, S6) analyzed data and played a major role in interpretation of data and preparation of the manuscript; DRA conceived of the project, helped with experimental design and analysis and interpretation of data and wrote the paper with AK. All authors were involved in discussing and interpreting data.

## Figure legends

**Figure S1:**
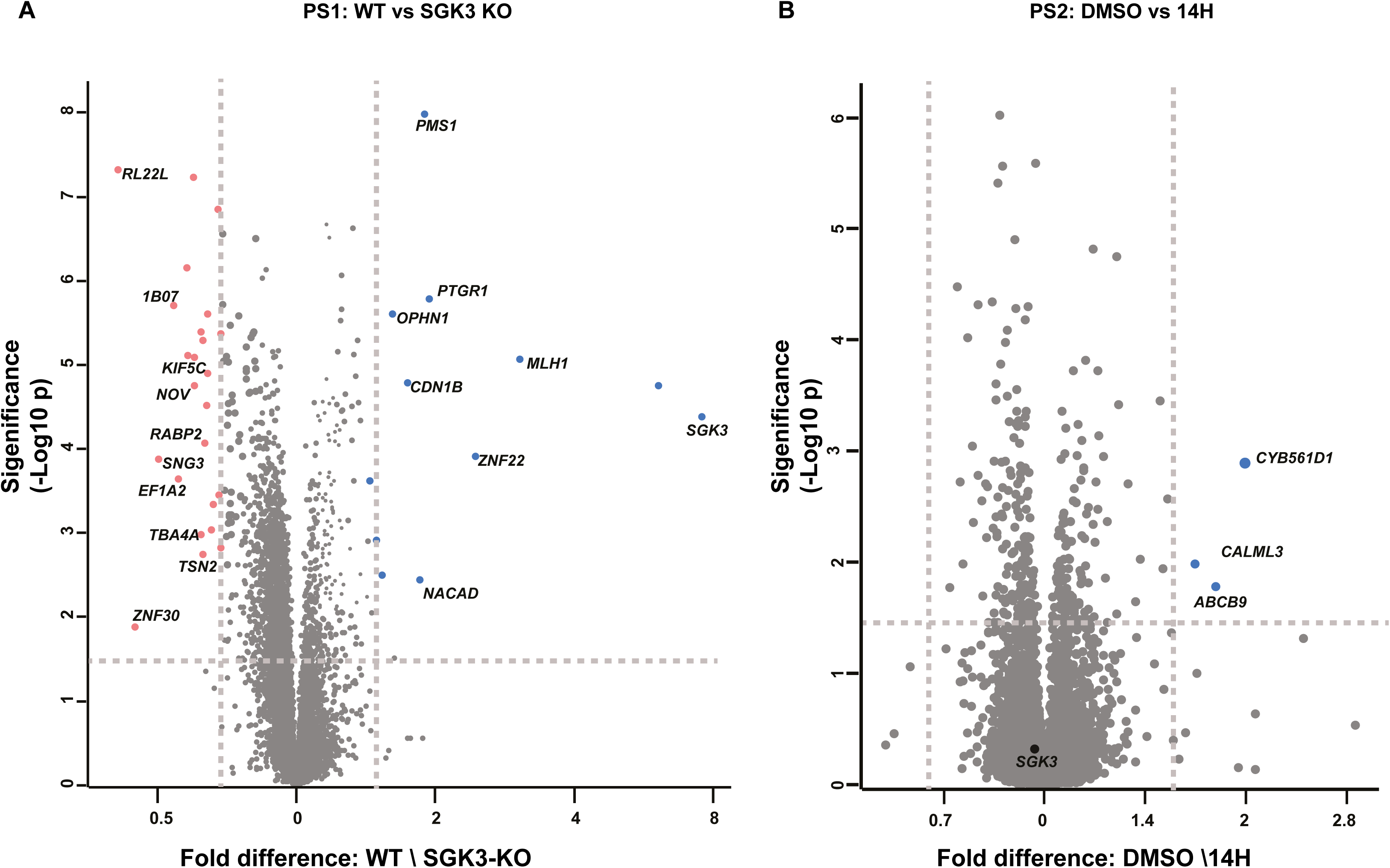
Total proteome changes in the phosphoproteomic screens. Volcano plot depicting the total proteome statistical significance between the HEK 293 Wild type (WT) and SGK3 knock-out (KO) (N= 4) from PS1 on the left panel and for the HEK293 cells treated DMSO and 14H conditions (N=3) from PS2 on the right panel. The significantly enriched protein groups in WT HEK293 cells are highlighted in blue filled circles and protein groups that are enriched in KO cells are highlighted in red filled circles. For PS2, the significantly enriched protein groups in DMSO treated cells are highlighted in blue filled circles and protein groups that are enriched in 14H treated cells are highlighted in red filled circles. The statistical significance was defined by the independent two-tailed t-test, which is corrected by permutation-based FDR of 5% using the Perseus software.

**Figure S2:**
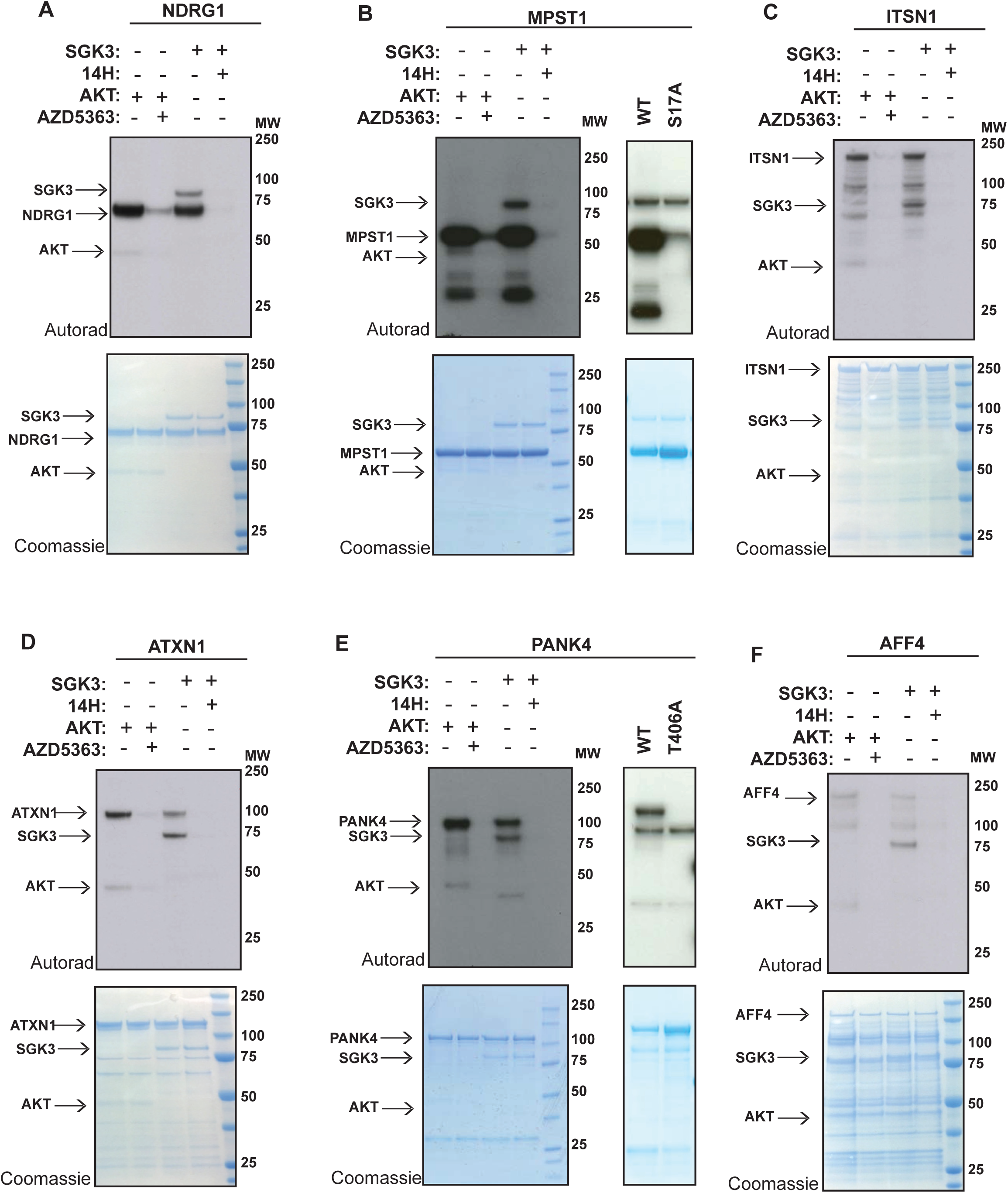
MPST1, PANK4, ITSN1, ATXN1 and AFF4 are phosphorylated by both SGK3 and Akt *in vitro*. 1-2 µg of GST-NDRG1 (A), GST-MPST (B), GST-ITSN1 (C), GST-ATXN1 (D), GST-PANK4 (E) and GST-AFF4 (F), were incubated with 0.5 µg active SGK3 or 0.125 µg active Akt and [γ^32^P]-ATP, in the presence or absence of 14H or AZD5363 (1 µM) for 30 min. The reactions were terminated by the addition of SDS loading buffer and separated by SDS/PAGE. Incorporation of [γ^32^P]-ATP was detected by autoradiography (top panels) and proteins were detected by Coomassie staining (bottom panels).

**Figure S3:**
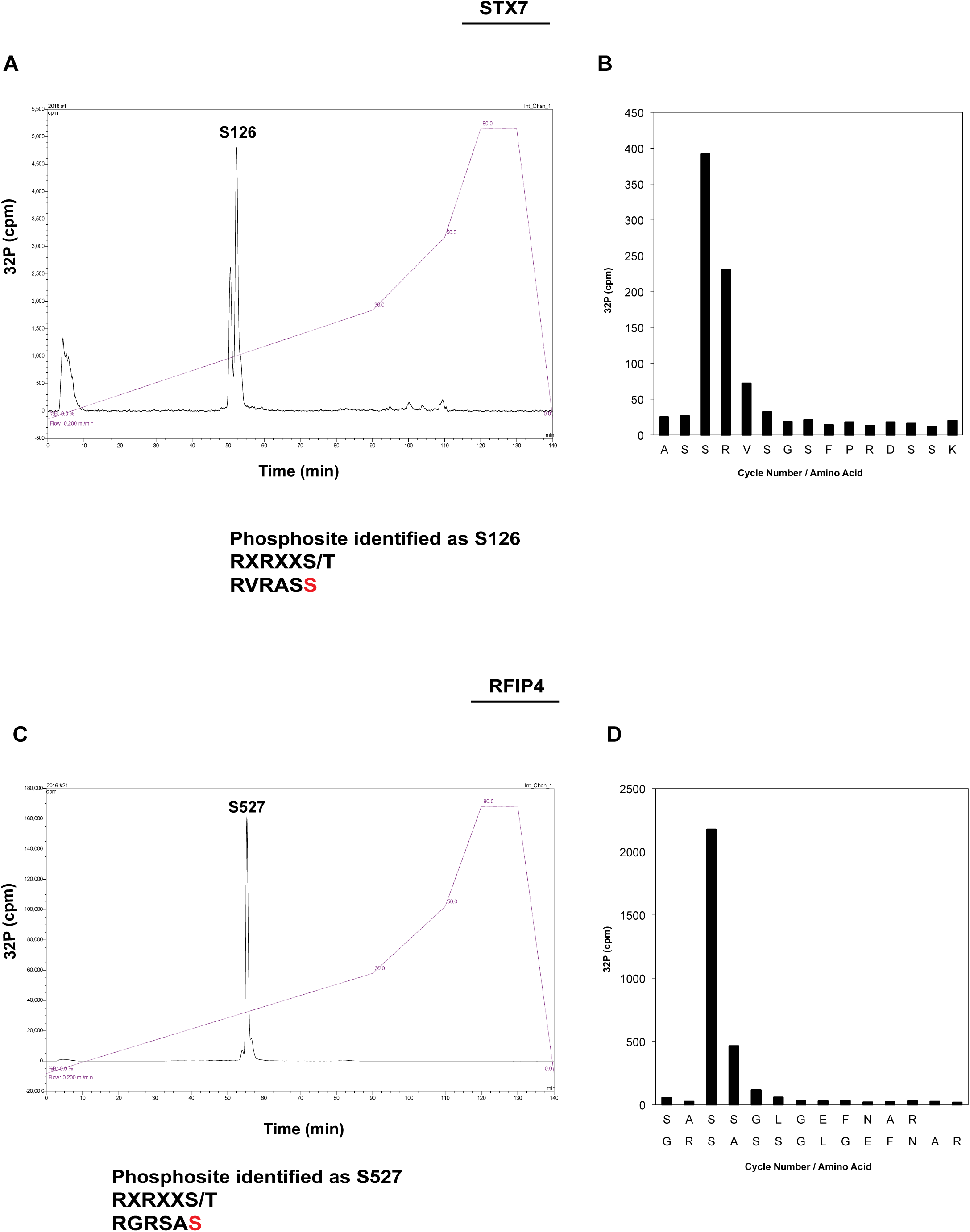
Phosphomapping of STX7 and RFIP4 by Edman degradation and mass spectrometry. GST-STX7 (A, B) and GST-RFIP4 (C,D) were incubated with active SGK3 in the presence of [γ^32^P]-ATP for 80 min. The reactions were terminated by the addition of SDS loading buffer and separated by SDS/PAGE. Proteins were detected by Coomassie staining and the bands corresponding to GST-SXT7 or GST-RFIP4 were excised from the gel and subjected to trypsin digestion. The resultant peptides were separated by HPLC (A, GST-STX7; C, GST-RFIP4) and ^32^P radioactivity was detected using an online radioactivity detector. The largest peak fractions were analyzed by Edman degradation and mass spectrometry (B, GST-STX7; D, GST-RFIP4).

**Figure S4:**
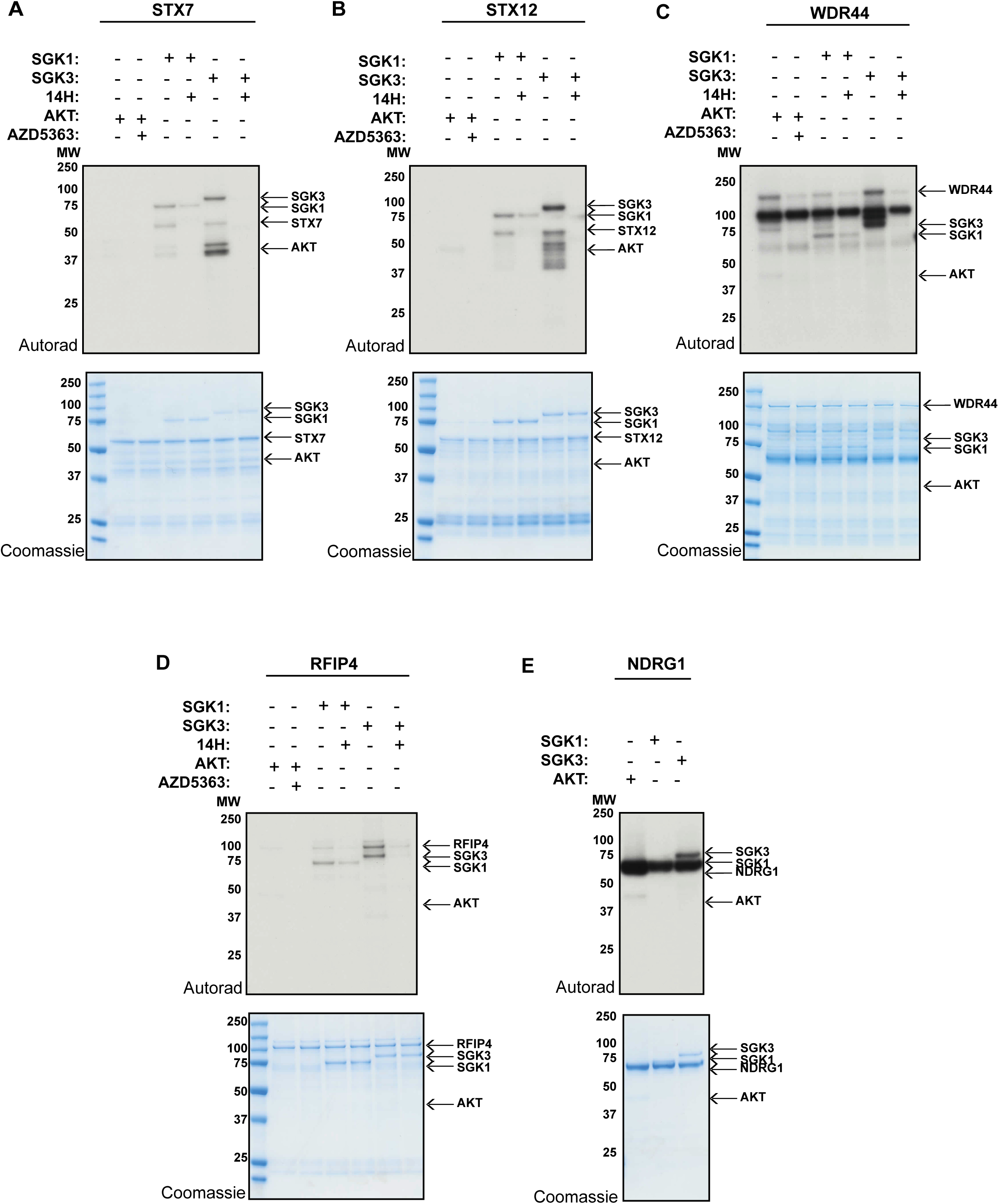
SGK1 can phosphorylate STX7, STX12, RFIP4 and WDR44 *in vitro*. 1-2 µg of GST-STX7 (A), GST-STX12 (B), GST-WDR44 (C) and GST-RFIP4 (D) were incubated with 0.5 µg active SGK1, 0.25 µg active SGK3 or 0.08 µg active Akt and [γ^32^P]-ATP, in the presence or absence of 14H or AZD5363 (1 µM) for 30 min. The reactions were terminated by the addition of SDS loading buffer and separated by SDS/PAGE. Proteins were detected by Coomassie staining (bottom panels) and incorporation of [γ^32^P]-ATP was detected by autoradiography (top panels).

**Figure S5:**
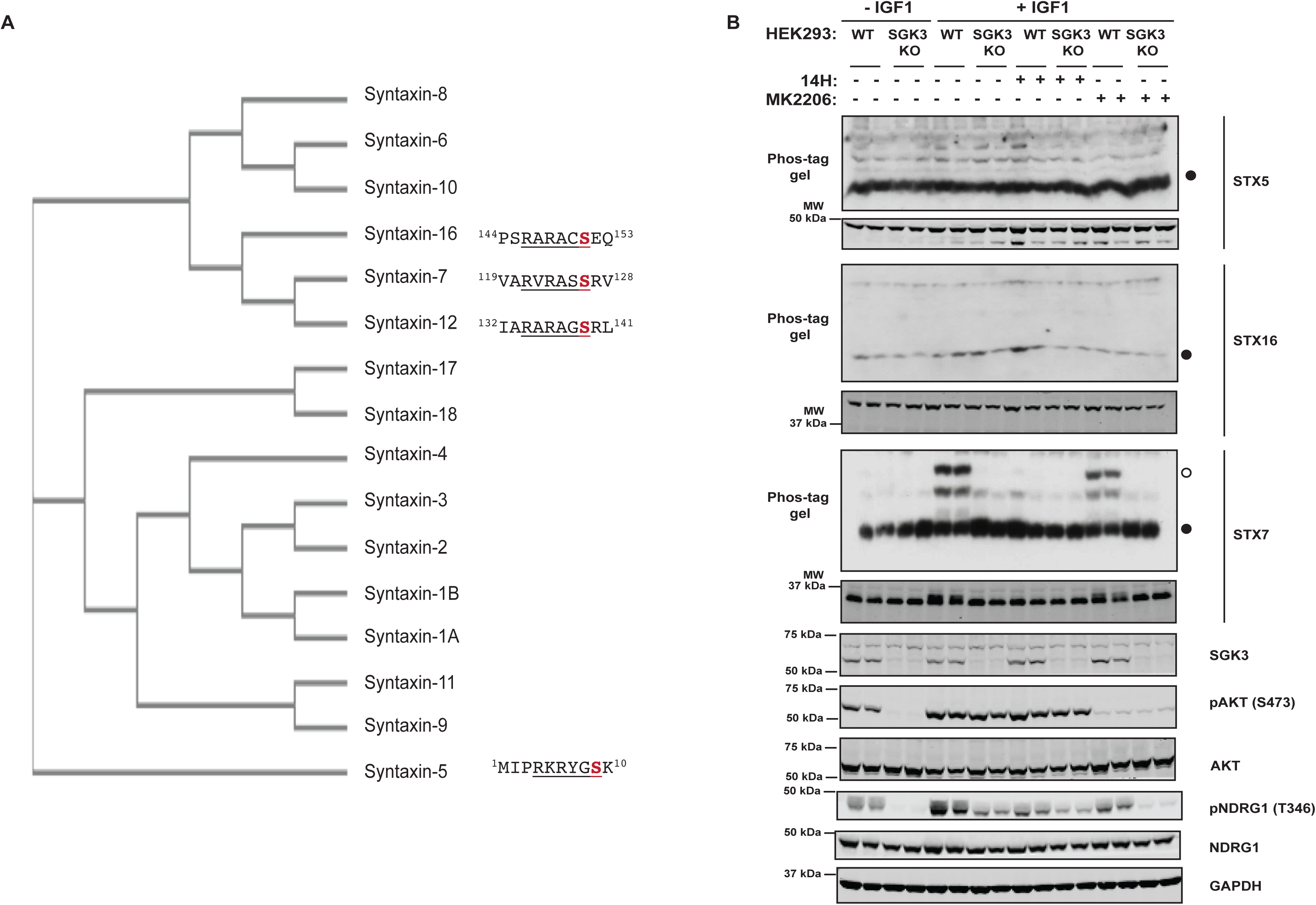
STX5 and STX16 are not phosphorylated by SGK3 in HEK293 cells. (A) Phylogenetic analysis of the Syntaxins. Sequence alignment and phylogenetic analysis were performed using Clustal Omega. The syntaxins containing a potential SGK3 phosphomotif are indicated. (B) Wild type (WT) and SGK3 knock-out (KO) HEK293 cells were serum starved overnight, then treated for 1 h with 14H (1 µM), MK2206 (1 µM) or with DMSO, followed by stimulation with IGF1 (50 ng/ml) for 15 min. STX5 and STX16 phosphorylation was analyzed by Phos-tag assays (top panels, top panels, °□phosphorylated, ^•^□non-phosphorylated,). Phos-tag analysis of the same samples for STX7 was performed as a control. Control immunoblots were performed on normal gels with the indicated antibodies. Immunoblots were developed using the LI-COR Odyssey CLx Western Blot imaging system analysis with the indicated antibodies at 0.5-1 µg/mL concentration.

**Figure S6:**
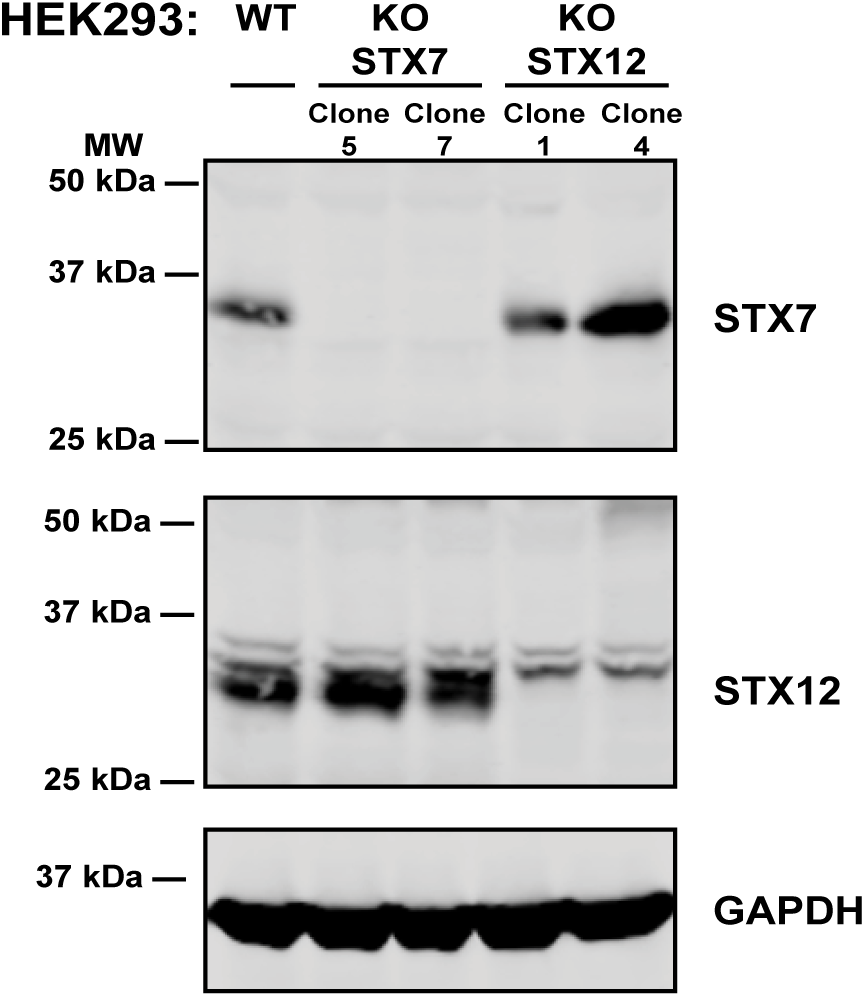
Generation of STX7 and STX12 knock-out HEK293 cell lines by CRISPR/Cas9. For generating the STX7 knock-out (KO) cell line the guide RNAs targeted exon 2 of *STX7* gene, while for STX12 KO the guide RNAs targeted exon 1 of *STX12* gene. Two representative KO clones for each cell line are shown. All the clones were blotted for STX7, STX12 and GAPDH as a control.

## References

1. Fruman, D. A., Chiu, H., Hopkins, B. D., Bagrodia, S., Cantley, L. C. and Abraham, R. T. (2017) The PI3K Pathway in Human Disease. Cell. 170, 605–635

2. Okkenhaug, K., Graupera, M. and Vanhaesebroeck, B. (2016) Targeting PI3K in Cancer: Impact on Tumor Cells, Their Protective Stroma, Angiogenesis, and Immunotherapy. Cancer Discov. 6, 1090–1105

3. Andre, F., Ciruelos, E., Rubovszky, G., Campone, M., Loibl, S., Rugo, H. S., Iwata, H., Conte, P., Mayer, I. A., Kaufman, B., Yamashita, T., Lu, Y. S., Inoue, K., Takahashi, M., Papai, Z., Longin, A. S., Mills, D., Wilke, C., Hirawat, S., Juric, D. and Group, S.-S. (2019) Alpelisib for PIK3CA-Mutated, Hormone Receptor-Positive Advanced Breast Cancer. N Engl J Med. 380, 1929–1940

4. Turner, N. C., Alarcon, E., Armstrong, A. C., Philco, M., Lopez Chuken, Y. A., Sablin, M. P., Tamura, K., Gomez Villanueva, A., Perez-Fidalgo, J. A., Cheung, S. Y. A., Corcoran, C., Cullberg, M., Davies, B. R., de Bruin, E. C., Foxley, A., Lindemann, J. P. O., Maudsley, R., Moschetta, M., Outhwaite, E., Pass, M., Rugman, P., Schiavon, G. and Oliveira, M. (2019) BEECH: a dose-finding run-in followed by a randomised phase II study assessing the efficacy of AKT inhibitor capivasertib (AZD5363) combined with paclitaxel in patients with estrogen receptor-positive advanced or metastatic breast cancer, and in a PIK3CA mutant sub-population. Ann Oncol. 30, 774–780

5. Oliveira, M., Saura, C., Nuciforo, P., Calvo, I., Andersen, J., Passos-Coelho, J. L., Gil Gil, M., Bermejo, B., Patt, D. A., Ciruelos, E., de la Pena, L., Xu, N., Wongchenko, M., Shi, Z., Singel, S. M. and Isakoff, S. J. (2019) FAIRLANE, a double-blind placebo-controlled randomized phase II trial of neoadjuvant ipatasertib plus paclitaxel for early triple-negative breast cancer. Ann Oncol

6. Klempner, S. J., Myers, A. P. and Cantley, L. C. (2013) What a tangled web we weave: emerging resistance mechanisms to inhibition of the phosphoinositide 3-kinase pathway. Cancer Discov. 3, 1345–1354

7. Castel, P. and Scaltriti, M. (2018) Mechanisms of Resistance to PI3K and AKT Inhibitors. In Resistance to Targeted Anti-Cancer Therapeutics book series (RTACT, volume 15) (Bonavida, B., ed.). pp. 117–146, Springer

8. Tessier, M. and Woodgett, J. R. (2006) Serum and glucocorticoid-regulated protein kinases: variations on a theme. J Cell Biochem. 98, 1391–1407

9. Pearce, L. R., Komander, D. and Alessi, D. R. (2010) The nuts and bolts of AGC protein kinases. Nat Rev Mol Cell Biol. 11, 9–22

10. Sommer, E. M., Dry, H., Cross, D., Guichard, S., Davies, B. R. and Alessi, D. R. (2013) Elevated SGK1 predicts resistance of breast cancer cells to Akt inhibitors. Biochem J. 452, 499–508

11. Castel, P., Ellis, H., Bago, R., Toska, E., Razavi, P., Carmona, F. J., Kannan, S., Verma, C. S., Dickler, M., Chandarlapaty, S., Brogi, E., Alessi, D. R., Baselga, J. and Scaltriti, M. (2016) PDK1-SGK1 Signaling Sustains AKT-Independent mTORC1 Activation and Confers Resistance to PI3Kalpha Inhibition. Cancer Cell. 30, 229–242

12. Bago, R., Sommer, E., Castel, P., Crafter, C., Bailey, F. P., Shpiro, N., Baselga, J., Cross, D., Eyers, P. A. and Alessi, D. R. (2016) The hVps34-SGK3 pathway alleviates sustained PI3K/Akt inhibition by stimulating mTORC1 and tumour growth. Embo J. 35, 1902–1922

13. Kobayashi, T., Deak, M., Morrice, N. and Cohen, P. (1999) Characterization of the structure and regulation of two novel isoforms of serum- and glucocorticoid-induced protein kinase. Biochem J. 344, 189–197

14. Kobayashi, T. and Cohen, P. (1999) Activation of serum- and glucocorticoid-regulated protein kinase by agonists that activate phosphatidylinositide 3-kinase is mediated by 3-phosphoinositide-dependent protein kinase-1 (PDK1) and PDK2. Biochem J. 339, 319–328

15. Garcia-Martinez, J. M. and Alessi, D. R. (2008) mTOR complex 2 (mTORC2) controls hydrophobic motif phosphorylation and activation of serum- and glucocorticoid-induced protein kinase 1 (SGK1). Biochem J. 416, 375–385

16. Pearce, L. R., Sommer, E. M., Sakamoto, K., Wullschleger, S. and Alessi, D. R. (2011) Protor-1 is required for efficient mTORC2-mediated activation of SGK1 in the kidney. Biochem J. 436, 169–179

17. Bago, R., Malik, N., Munson, M. J., Prescott, A. R., Davies, P., Sommer, E., Shpiro, N., Ward, R., Cross, D., Ganley, I. G. and Alessi, D. R. (2014) Characterization of VPS34-IN1, a selective inhibitor of Vps34, reveals that the phosphatidylinositol 3-phosphate-binding SGK3 protein kinase is a downstream target of class III phosphoinositide 3-kinase. Biochem J. 463, 413–427

18. Biondi, R. M., Kieloch, A., Currie, R. A., Deak, M. and Alessi, D. R. (2001) The PIF-binding pocket in PDK1 is essential for activation of S6K and SGK, but not PKB. Embo J. 20, 4380–4390.

19. Tessier, M. and Woodgett, J. R. (2006) Role of the Phox homology domain and phosphorylation in activation of serum and glucocorticoid-regulated kinase-3. J Biol Chem. 281, 23978–23989

20. Virbasius, J. V., Song, X., Pomerleau, D. P., Zhan, Y., Zhou, G. W. and Czech, M. P. (2001) Activation of the Akt-related cytokine-independent survival kinase requires interaction of its phox domain with endosomal phosphatidylinositol 3-phosphate. Proc Natl Acad Sci U S A. 98, 12908–12913.

21. Gasser, J. A., Inuzuka, H., Lau, A. W., Wei, W., Beroukhim, R. and Toker, A. (2014) SGK3 mediates INPP4B-dependent PI3K signaling in breast cancer. Mol Cell. 56, 595–607

22. Malik, N., Macartney, T., Hornberger, A., Anderson, K. E., Tovell, H., Prescott, A. R. and Alessi, D. R. (2018) Mechanism of activation of SGK3 by growth factors via the Class 1 and Class 3 PI3Ks. Biochem J. 475, 117–135

23. Alessi, D. R., Caudwell, F. B., Andjelkovic, M., Hemmings, B. A. and Cohen, P. (1996) Molecular basis for the substrate specificity of protein kinase B; comparison with MAPKAP kinase-1 and p70 S6 kinase. FEBS letters. 399, 333–338

24. Obata, T., Yaffe, M. B., Leparc, G. G., Piro, E. T., Maegawa, H., Kashiwagi, A., Kikkawa, R. and Cantley, L. C. (2000) Peptide and Protein Library Screening Defines Optimal Substrate Motifs for AKT/PKB. J Biol Chem. 275, 36108–36115

25. Liu, D., Yang, X. and Songyang, Z. (2000) Identification of CISK, a new member of the SGK kinase family that promotes IL-3-dependent survival. Curr Biol. 10, 1233–1236

26. Snyder, P. M., Olson, D. R. and Thomas, B. C. (2002) Serum and glucocorticoid-regulated kinase modulates Nedd4-2-mediated inhibition of the epithelial Na+ channel. J Biol Chem. 277, 5–8

27. Bhalla, V., Daidie, D., Li, H., Pao, A. C., LaGrange, L. P., Wang, J., Vandewalle, A., Stockand, J. D., Staub, O. and Pearce, D. (2005) Serum- and glucocorticoid-regulated kinase 1 regulates ubiquitin ligase neural precursor cell-expressed, developmentally down-regulated protein 4-2 by inducing interaction with 14-3-3. Mol Endocrinol. 19, 3073–3084

28. Lee, I. H., Dinudom, A., Sanchez-Perez, A., Kumar, S. and Cook, D. I. (2007) Akt mediates the effect of insulin on epithelial sodium channels by inhibiting Nedd4-2. J Biol Chem. 282, 29866–29873

29. Brunet, A., Park, J., Tran, H., Hu, L. S., Hemmings, B. A. and Greenberg, M. E. (2001) Protein kinase SGK mediates survival signals by phosphorylating the forkhead transcription factor FKHRL1 (FOXO3a). Mol Cell Biol. 21, 952–965

30. Murray, J. T., Campbell, D. G., Morrice, N., Auld, G. C., Shpiro, N., Marquez, R., Peggie, M., Bain, J., Bloomberg, G. B., Grahammer, F., Lang, F., Wulff, P., Kuhl, D. and Cohen, P. (2004) Exploitation of KESTREL to identify N-myc downstream-regulated gene family members as physiological substrates for SGK1 and GSK3. Biochem J

31. Slagsvold, T., Marchese, A., Brech, A. and Stenmark, H. (2006) CISK attenuates degradation of the chemokine receptor CXCR4 via the ubiquitin ligase AIP4. Embo J. 25, 3738–3749

32. Xu, J., Liao, L., Qin, J., Xu, J., Liu, D. and Songyang, Z. (2009) Identification of Flightless-I as a substrate of the cytokine-independent survival kinase CISK. J Biol Chem. 284, 14377–14385

33. Murray, J. T., Cummings, L. A., Bloomberg, G. B. and Cohen, P. (2005) Identification of different specificity requirements between SGK1 and PKBalpha. FEBS letters. 579, 991–994

34. Erickson, B. K., Jedrychowski, M. P., McAlister, G. C., Everley, R. A., Kunz, R. and Gygi, S. P. (2015) Evaluating multiplexed quantitative phosphopeptide analysis on a hybrid quadrupole mass filter/linear ion trap/orbitrap mass spectrometer. Anal Chem. 87, 1241–1249

35. Cox, J. and Mann, M. (2011) Quantitative, high-resolution proteomics for data-driven systems biology. Annual review of biochemistry. 80, 273–299

36. Humphrey, S. J., Azimifar, S. B. and Mann, M. (2015) High-throughput phosphoproteomics reveals in vivo insulin signaling dynamics. Nat Biotechnol. 33, 990–995

37. Vizcaino, J. A., Csordas, A., del-Toro, N., Dianes, J. A., Griss, J., Lavidas, I., Mayer, G., Perez-Riverol, Y., Reisinger, F., Ternent, T., Xu, Q. W., Wang, R. and Hermjakob, H. (2016) 2016 update of the PRIDE database and its related tools. Nucleic Acids Res. 44, D447–456

38. Brown, K. K., Montaser-Kouhsari, L., Beck, A. H. and Toker, A. (2015) MERIT40 Is an Akt Substrate that Promotes Resolution of DNA Damage Induced by Chemotherapy. Cell reports. 11, 1358–1366

39. Chen, H. K., Fernandez-Funez, P., Acevedo, S. F., Lam, Y. C., Kaytor, M. D., Fernandez, M. H., Aitken, A., Skoulakis, E. M., Orr, H. T., Botas, J. and Zoghbi, H. Y. (2003) Interaction of Akt-phosphorylated ataxin-1 with 14-3-3 mediates neurodegeneration in spinocerebellar ataxia type 1. Cell. 113, 457–468

40. Walia, V., Cuenca, A., Vetter, M., Insinna, C., Perera, S., Lu, Q., Ritt, D. A., Semler, E., Specht, S., Stauffer, J., Morrison, D. K., Lorentzen, E. and Westlake, C. J. (2019) Akt Regulates a Rab11-Effector Switch Required for Ciliogenesis. Developmental cell. 50, 229–246 e227

41. Jenna, S., Hussain, N. K., Danek, E. I., Triki, I., Wasiak, S., McPherson, P. S. and Lamarche-Vane, N. (2002) The activity of the GTPase-activating protein CdGAP is regulated by the endocytic protein intersectin. J Biol Chem. 277, 6366–6373

42. Mullock, B. M., Smith, C. W., Ihrke, G., Bright, N. A., Lindsay, M., Parkinson, E. J., Brooks, D. A., Parton, R. G., James, D. E., Luzio, J. P. and Piper, R. C. (2000) Syntaxin 7 is localized to late endosome compartments, associates with Vamp 8, and Is required for late endosome-lysosome fusion. Mol Biol Cell. 11, 3137–3153

43. Tang, B. L., Tan, A. E., Lim, L. K., Lee, S. S., Low, D. Y. and Hong, W. (1998) Syntaxin 12, a member of the syntaxin family localized to the endosome. J Biol Chem. 273, 6944–6950

44. Wallace, D. M., Lindsay, A. J., Hendrick, A. G. and McCaffrey, M. W. (2002) The novel Rab11-FIP/Rip/RCP family of proteins displays extensive homo- and hetero-interacting abilities. Biochem Biophys Res Commun. 292, 909–915

45. Zeng, J., Ren, M., Gravotta, D., De Lemos-Chiarandini, C., Lui, M., Erdjument-Bromage, H., Tempst, P., Xu, G., Shen, T. H., Morimoto, T., Adesnik, M. and Sabatini, D. D. (1999) Identification of a putative effector protein for rab11 that participates in transferrin recycling. Proc Natl Acad Sci U S A. 96, 2840–2845

46. Kinoshita, E., Kinoshita-Kikuta, E., Takiyama, K. and Koike, T. (2006) Phosphate-binding tag, a new tool to visualize phosphorylated proteins. Mol Cell Proteomics. 5, 749–757

47. Ito, G., Katsemonova, K., Tonelli, F., Lis, P., Baptista, M. A., Shpiro, N., Duddy, G., Wilson, S., Ho, P. W., Ho, S. L., Reith, A. D. and Alessi, D. R. (2016) Phos-tag analysis of Rab10 phosphorylation by LRRK2: a powerful assay for assessing kinase function and inhibitors. Biochem J. 473, 2671–2685

48. Najafov, A., Sommer, E. M., Axten, J. M., Deyoung, M. P. and Alessi, D. R. (2011) Characterization of GSK2334470, a novel and highly specific inhibitor of PDK1. Biochem J. 433, 357–369

49. Chresta, C. M., Davies, B. R., Hickson, I., Harding, T., Cosulich, S., Critchlow, S. E., Vincent, J. P., Ellston, R., Jones, D., Sini, P., James, D., Howard, Z., Dudley, P., Hughes, G., Smith, L., Maguire, S., Hummersone, M., Malagu, K., Menear, K., Jenkins, R., Jacobsen, M., Smith, G. C., Guichard, S. and Pass, M. (2010) AZD8055 is a potent, selective, and orally bioavailable ATP-competitive mammalian target of rapamycin kinase inhibitor with in vitro and in vivo antitumor activity. Cancer Res. 70, 288–298

50. Folkes, A. J., Ahmadi, K., Alderton, W. K., Alix, S., Baker, S. J., Box, G., Chuckowree, I. S., Clarke, P. A., Depledge, P., Eccles, S. A., Friedman, L. S., Hayes, A., Hancox, T. C., Kugendradas, A., Lensun, L., Moore, P., Olivero, A. G., Pang, J., Patel, S., Pergl-Wilson, G. H., Raynaud, F. I., Robson, A., Saghir, N., Salphati, L., Sohal, S., Ultsch, M. H., Valenti, M., Wallweber, H. J., Wan, N. C., Wiesmann, C., Workman, P., Zhyvoloup, A., Zvelebil, M. J. and Shuttleworth, S. J. (2008) The identification of 2-(1H-indazol-4-yl)-6-(4-methanesulfonyl-piperazin-1-ylmethyl)-4-morpholin-4-yl-thieno[3,2-d]pyrimidine (GDC-0941) as a potent, selective, orally bioavailable inhibitor of class I PI3 kinase for the treatment of cancer. J Med Chem. 51, 5522–5532

51. Brandhorst, D., Zwilling, D., Rizzoli, S. O., Lippert, U., Lang, T. and Jahn, R. (2006) Homotypic fusion of early endosomes: SNAREs do not determine fusion specificity. Proc Natl Acad Sci U S A. 103, 2701–2706

52. Zwilling, D., Cypionka, A., Pohl, W. H., Fasshauer, D., Walla, P. J., Wahl, M. C. and Jahn, R. (2007) Early endosomal SNAREs form a structurally conserved SNARE complex and fuse liposomes with multiple topologies. Embo J. 26, 9–18

53. Coleman, M. P. and Freeman, M. R. (2010) Wallerian degeneration, wld(s), and nmnat. Annual review of neuroscience. 33, 245–267

54. Xiong, X., Hao, Y., Sun, K., Li, J., Li, X., Mishra, B., Soppina, P., Wu, C., Hume, R. I. and Collins, C. A. (2012) The Highwire ubiquitin ligase promotes axonal degeneration by tuning levels of Nmnat protein. PLoS Biol. 10, e1001440

55. Pao, K. C., Wood, N. T., Knebel, A., Rafie, K., Stanley, M., Mabbitt, P. D., Sundaramoorthy, R., Hofmann, K., van Aalten, D. M. F. and Virdee, S. (2018) Activity-based E3 ligase profiling uncovers an E3 ligase with esterification activity. Nature. 556, 381–385

56. Garcia-Martinez, J. M., Moran, J., Clarke, R. G., Gray, A., Cosulich, S. C., Chresta, C. M. and Alessi, D. R. (2009) Ku-0063794 is a specific inhibitor of the mammalian target of rapamycin (mTOR). Biochem J. 421, 29–42

57. Jahn, R. and Scheller, R. H. (2006) SNAREs--engines for membrane fusion. Nat Rev Mol Cell Biol. 7, 631–643

58. Teng, F. Y., Wang, Y. and Tang, B. L. (2001) The syntaxins. Genome Biol. 2, REVIEWS3012

59. Trost, M., English, L., Lemieux, S., Courcelles, M., Desjardins, M. and Thibault, P. (2009) The phagosomal proteome in interferon-gamma-activated macrophages. Immunity. 30, 143–154

60. Trost, M., Bridon, G., Desjardins, M. and Thibault, P. (2010) Subcellular phosphoproteomics. Mass Spectrom Rev. 29, 962–990

61. Antonin, W., Riedel, D. and von Mollard, G. F. (2000) The SNARE Vti1a-beta is localized to small synaptic vesicles and participates in a novel SNARE complex. The Journal of neuroscience: the official journal of the Society for Neuroscience. 20, 5724–5732

62. Achuthan, A., Masendycz, P., Lopez, J. A., Nguyen, T., James, D. E., Sweet, M. J., Hamilton, J. A. and Scholz, G. M. (2008) Regulation of the endosomal SNARE protein syntaxin 7 by colony-stimulating factor 1 in macrophages. Mol Cell Biol. 28, 6149–6159

63. Jani, R. A., Purushothaman, L. K., Rani, S., Bergam, P. and Setty, S. R. (2015) STX13 regulates cargo delivery from recycling endosomes during melanosome biogenesis. J Cell Sci. 128, 3263–3276

64. Dulubova, I., Sugita, S., Hill, S., Hosaka, M., Fernandez, I., Sudhof, T. C. and Rizo, J. (1999) A conformational switch in syntaxin during exocytosis: role of munc18. Embo J. 18, 4372–4382

65. Munson, M., Chen, X., Cocina, A. E., Schultz, S. M. and Hughson, F. M. (2000) Interactions within the yeast t-SNARE Sso1p that control SNARE complex assembly. Nat Struct Biol. 7, 894–902

66. Antonin, W., Dulubova, I., Arac, D., Pabst, S., Plitzner, J., Rizo, J. and Jahn, R. (2002) The N-terminal domains of syntaxin 7 and vti1b form three-helix bundles that differ in their ability to regulate SNARE complex assembly. J Biol Chem. 277, 36449–36456

67. Foletti, D. L., Lin, R., Finley, M. A. and Scheller, R. H. (2000) Phosphorylated syntaxin 1 is localized to discrete domains along a subset of axons. The Journal of neuroscience: the official journal of the Society for Neuroscience. 20, 4535–4544

68. Chung, S. H., Polgar, J. and Reed, G. L. (2000) Protein kinase C phosphorylation of syntaxin 4 in thrombin-activated human platelets. J Biol Chem. 275, 25286–25291

69. Tian, J. H., Das, S. and Sheng, Z. H. (2003) Ca2+-dependent phosphorylation of syntaxin-1A by the death-associated protein (DAP) kinase regulates its interaction with Munc18. J Biol Chem. 278, 26265–26274

70. Liu, X., Heidelberger, R. and Janz, R. (2014) Phosphorylation of syntaxin 3B by CaMKII regulates the formation of t-SNARE complexes. Molecular and cellular neurosciences. 60, 53–62

71. Prekeris, R., Klumperman, J., Chen, Y. A. and Scheller, R. H. (1998) Syntaxin 13 mediates cycling of plasma membrane proteins via tubulovesicular recycling endosomes. J Cell Biol. 143, 957–971

72. Kean, M. J., Williams, K. C., Skalski, M., Myers, D., Burtnik, A., Foster, D. and Coppolino, M. G. (2009) VAMP3, syntaxin-13 and SNAP23 are involved in secretion of matrix metalloproteinases, degradation of the extracellular matrix and cell invasion. J Cell Sci. 122, 4089–4098

73. Larsen, M. R., Thingholm, T. E., Jensen, O. N., Roepstorff, P. and Jorgensen, T. J. (2005) Highly selective enrichment of phosphorylated peptides from peptide mixtures using titanium dioxide microcolumns. Mol Cell Proteomics. 4, 873–886

74. Hansen, M., Peltier, J., Killy, B., Amin, B., Bodendorfer, B., Hartlova, A., Uebel, S., Bosmann, M., Hofmann, J., Buttner, C., Ekici, A. B., Kuttke, M., Franzyk, H., Foged, C., Beer-Hammer, S., Schabbauer, G., Trost, M. and Lang, R. (2019) Macrophage Phosphoproteome Analysis Reveals MINCLE-dependent and -independent Mycobacterial Cord Factor Signaling. Mol Cell Proteomics. 18, 669–685

75. Munoz, I. M., Morgan, M. E., Peltier, J., Weiland, F., Gregorczyk, M., Brown, F. C., Macartney, T., Toth, R., Trost, M. and Rouse, J. (2018) Phosphoproteomic screening identifies physiological substrates of the CDKL5 kinase. Embo J. 37

76. Lai, Y. C., Kondapalli, C., Lehneck, R., Procter, J. B., Dill, B. D., Woodroof, H. I., Gourlay, R., Peggie, M., Macartney, T. J., Corti, O., Corvol, J. C., Campbell, D. G., Itzen, A., Trost, M. and Muqit, M. M. (2015) Phosphoproteomic screening identifies Rab GTPases as novel downstream targets of PINK1. Embo J. 34, 2840–2861

77. Cox, J. and Mann, M. (2008) MaxQuant enables high peptide identification rates, individualized p.p.b.-range mass accuracies and proteome-wide protein quantification. Nat Biotechnol. 26, 1367–1372

78. Tyanova, S., Temu, T. and Cox, J. (2016) The MaxQuant computational platform for mass spectrometry-based shotgun proteomics. Nat Protoc. 11, 2301–2319

79. Tyanova, S., Temu, T., Sinitcyn, P., Carlson, A., Hein, M. Y., Geiger, T., Mann, M. and Cox, J. (2016) The Perseus computational platform for comprehensive analysis of (prote)omics data. Nat Methods. 13, 731–740

